# PAX8 orchestrates an angiogenic program through interaction with SOX17

**DOI:** 10.1101/2020.09.09.290387

**Authors:** Daniele Chaves-Moreira, Marilyn A. Mitchell, Cristina Arruza, Priyanka Rawat, Simone Sidoli, Robbin Nameki, Jessica Reddy, Rosario I. Corona, Sisi Ma, Boris Winterhoff, Gottfried E. Konecny, Benjamin A. Garcia, Donita C. Brady, Kate Lawrenson, Patrice J. Morin, Ronny Drapkin

## Abstract

Worldwide, the number of new ovarian cancer cases approaches 300,000 with more than 180,000 deaths every year. The low survival-rate reflects the limitations of current therapies and highlights the importance of identifying new therapeutic targets. Despite significant recent efforts to identify novel vulnerabilities in ovarian cancer, none have led to effective durable therapies with improvement in overall survival. PAX8, a lineage-transcription factor, whose expression is a major molecular feature of ovarian carcinomas, represents a novel therapeutic target. Herein, we have identified SOX17 as a *bona fide* PAX8-interacting partner and elucidated the impact of this interaction on the development of ovarian cancer. Importantly, we found that PAX8 and SOX17 regulate tumor angiogenesis *in vitro* and *in vivo*. The role of PAX8 and SOX17 in the regulation of angiogenesis reveals a novel function for these factors in regulating the tumor microenvironment and highlight this pathway as a viable therapeutic target.

## INTRODUCTION

Epithelial ovarian carcinoma is the most lethal of all gynecologic malignancies, claiming an estimated 14,000 lives annually in the United States of America (1) with worldwide numbers approaching 180,000 deaths annually (2). The lack of effective screening tools result in the majority of cases being diagnosed at an advanced stage and thus translating into a 5-year survival rate of less than 30% (3). The absence of adequate screening is compounded by the propensity of this disease to acquire chemoresistance and relapse of disease in the majority of patients despite initial response to platinum-based chemotherapy and surgical cytoreduction. Although much progress in ovarian cancer disease knowledge has emerged, effective targeted therapies have yet to impact overall survival rates (4).

The knowledge of the pathogenesis of high-grade serous ovarian carcinoma (HGSOC), the most common subtype of this disease, has greatly advanced over the past decade. A number of studies support the fallopian tube secretory epithelial cells (FTSEC) as the site of origin of the majority of HGSOC (5–13). The development of female reproductive tract is governed by the PAX8 transcription factor (14), which is a member of the Paired-Box family of transcription factors that play essential roles during embryogenesis and tumorigenesis (15). In normal oviducts, PAX8 expression is restricted to the secretory cells while neighboring ciliated cells show no expression. The sustained expression of PAX8 in the adult FTSEC and in nearly all HGSOCs (16), previously led us to use the PAX8 promoter to develop a genetically engineered mouse model of HGSOC (17, 18). Knockdown of PAX8 in ovarian cancer cells leads to apoptosis (19–21) supporting a definitive role for PAX8 in ovarian cancer growth and progression. However, it remains unclear how PAX8 drives the development of the Mullerian reproductive tract or how it supports neoplastic growth.

Surprisingly, PAX8 knockdown following transcriptome analysis revealed a negligible impact on the FTSEC cell lines with very few transcripts significantly affected by PAX8 loss (21, 22). In contrast, the range of transcripts altered by PAX8 loss in cancer cell lines was considerably higher. Ontology analysis of the alterations after PAX8 loss showed changes in proliferation, angiogenesis and adhesion pathways that are crucial for tumor progression (22). Chromatin immunoprecipitation-sequencing (ChIP-seq analysis) showed that the PAX8 cistrome is reprogrammed during the process of malignant transformation by the widespread redistribution of PAX8 binding sites in the genome of ovarian cancer cells. Moreover, a recent study showed that non-coding somatic mutations disrupt the PAX8 transcriptional program in ovarian cancer (23).

To further survey the roles of PAX8 in ovarian carcinomas, we purified the PAX8 protein complex from a panel of different fallopian tube secretory cells, and different ovarian carcinoma cells and identified its components. Our analyses reveal that PAX8 is part of a NuRD chromatin remodeling complex and associates with multiple factors including SOX17. We show that depletion of PAX8 or SOX17 impacts the expression of factors involved in angiogenesis and functionally disrupts capillary formation *in vitro* and in mouse models. Disruption in angiogenesis greatly decreased tumor burden, ascites formation and lead to improved survival. These findings support a role of PAX8 and SOX17 in the regulation of angiogenesis and highlight this pathway as a viable therapeutic target.

## RESULTS

### Biochemical purification of the PAX8 complex

To better understand the function of PAX8 in benign compared to malignant fallopian tube secretory cells, we developed a biochemical affinity-purification method (Figure 1A). First, we generated nuclear extracts as previously described (24) and purified the endogenous PAX8 protein complex from three ovarian carcinoma cell lines (OVCAR4, KURAMOCHI, and OVSAHO) and three immortalized fallopian tube secretory cell lines (FT194, FT246, and FT282) using anti-PAX8-specific antibodies. Immunoblotting following affinity chromatography demonstrated the specific enrichment of PAX8 in our system (Figure 1B). When the affinity-purified PAX8 complex was evaluated on size-exclusion chromatography it revealed a size of approximately 600 kDa (Figure 1C). Mass spectrometry analysis of the PAX8-containing fractions identified a number of putative PAX8-interacting proteins (Figure 1D and Table 1).

**Table 1:**
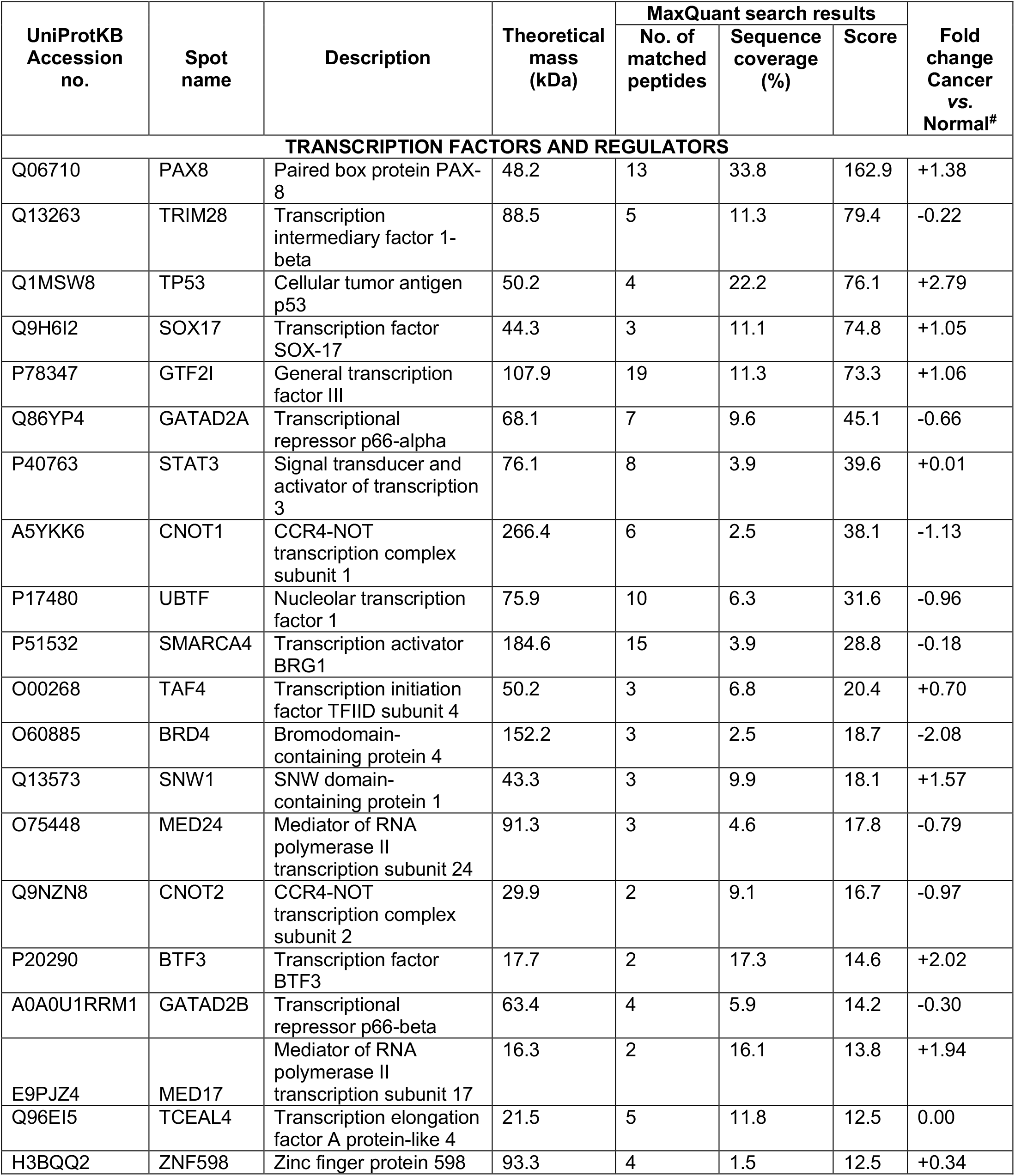
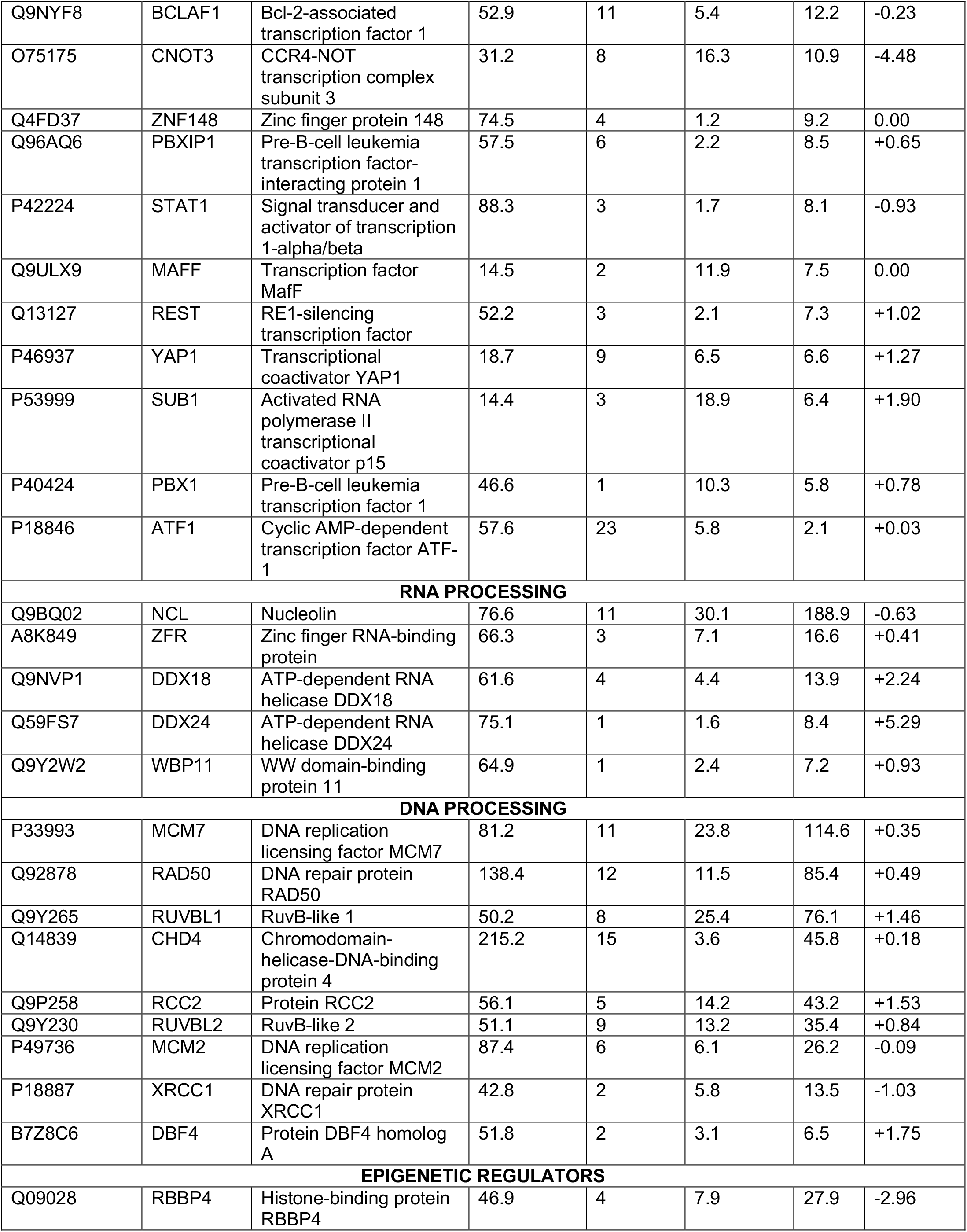
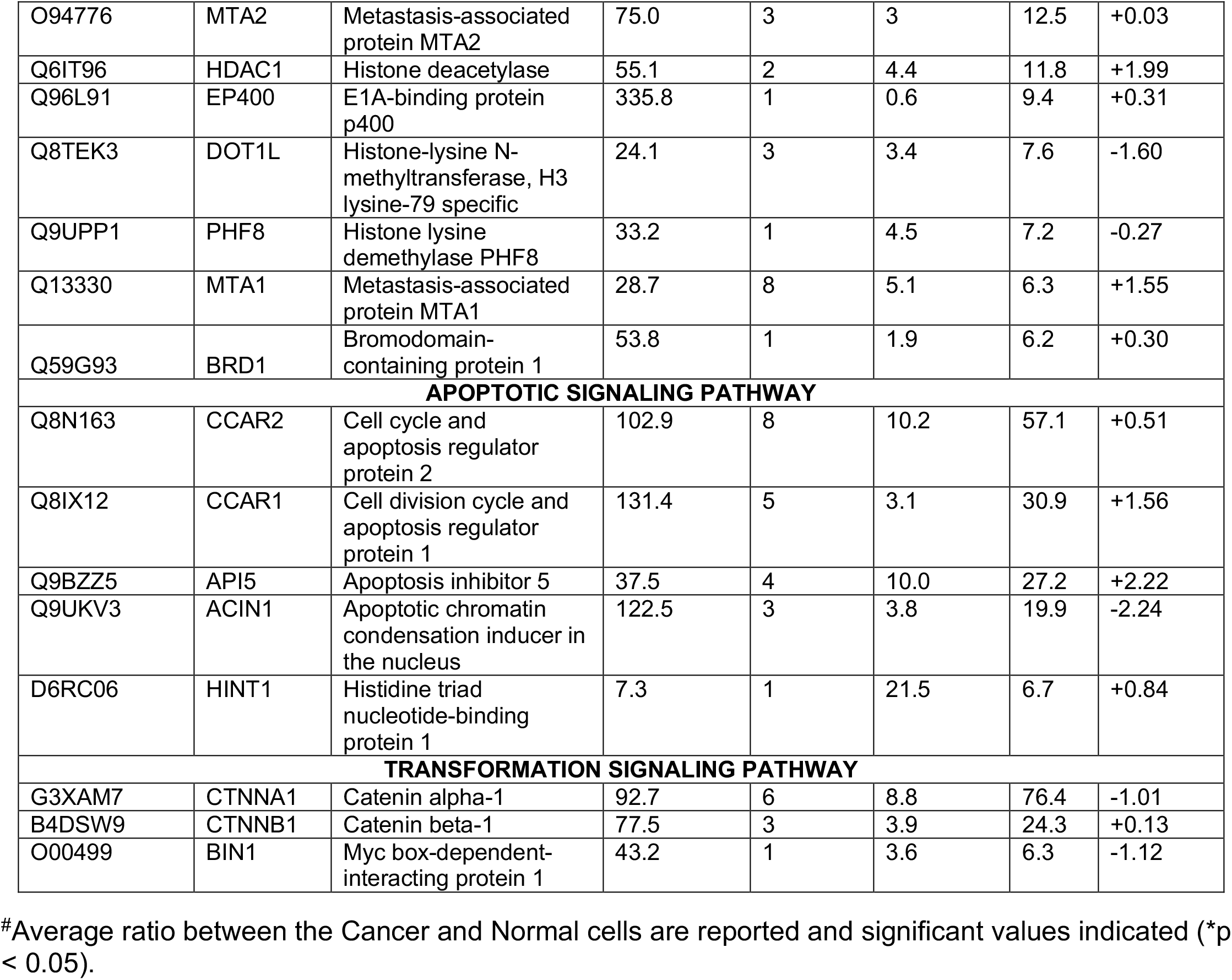
Putative PAX8-interacting partners.

**Figure 1:**
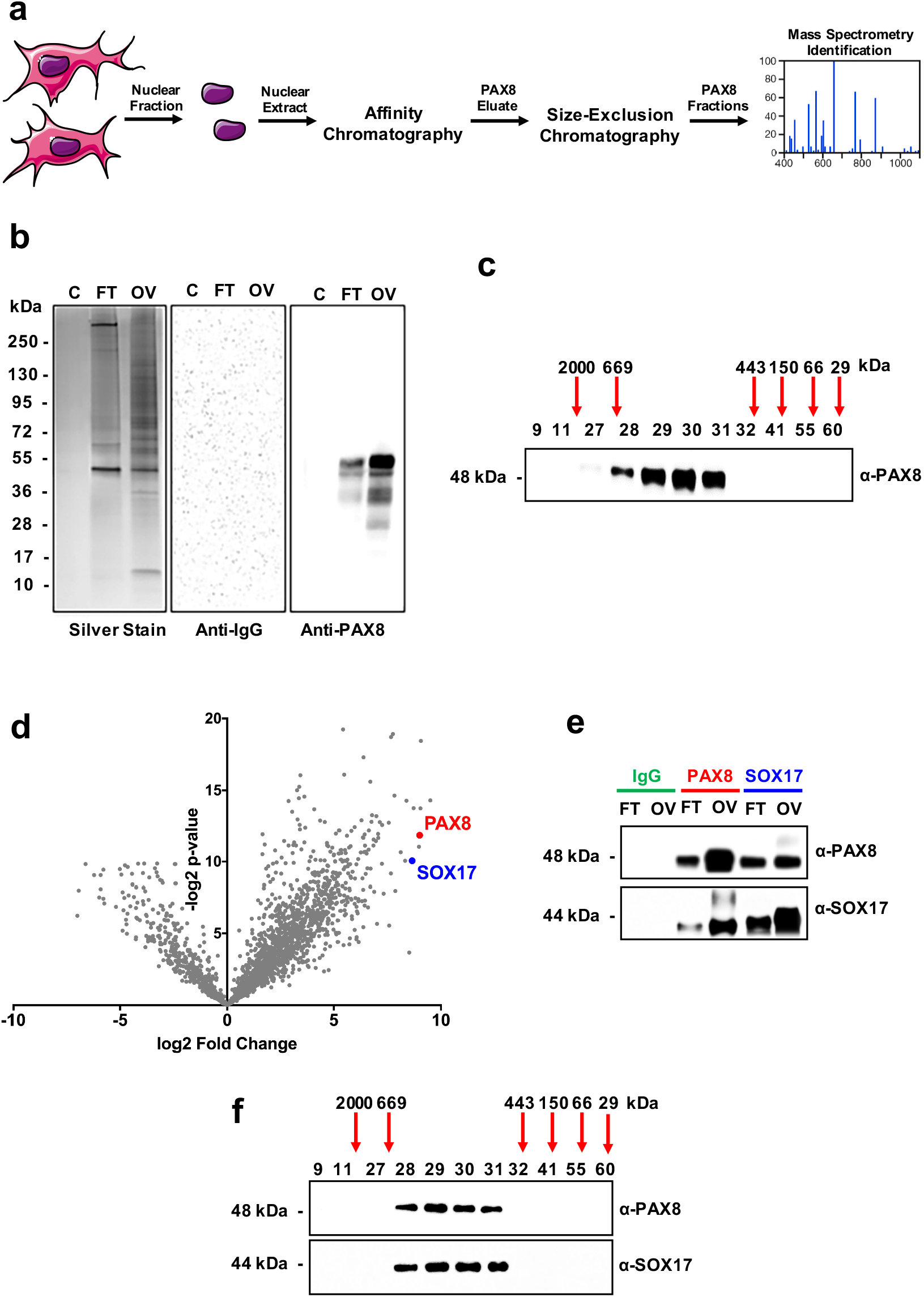
Three-step proteomic approach identifies putative PAX8-interacting partners. (a) Schematic of the workflow for the PAX8-interacting partner’s identification. (b) Representative examples of endogenous PAX8 immunoprecipitation silver-stained gel and immunoblot from fallopian tube secretory cells (FT) and ovarian carcinoma cells (OV). IMR90 cells, which are PAX8 negative, were used as negative control (C). (c) Immunoblot of size-exclusion fractions showing the presence of PAX8 at 600 kDa size range. PAX8 immunoprecipitates and gel-filtration fractions were sent for mass spectrometry analysis. (d) Volcano plot of identified putative PAX8-interacting partners. (e) Confirmation of PAX8 and SOX17 interaction by co-immunoprecipitation experiments from FTSEC and HGSOC cell lines. (f) PAX8 and SOX17 immunoblots of size exclusion fractions demonstrate co-elution at 600 kDa size range fractions.

Remarkably, many of the putative PAX8-interacting proteins represent components of chromatin-remodeling complexes. For example, chromodomain-helicase-DNA-binding protein 4 (CHD4), Transcriptional repressor p66-alpha (GATAD2A), Metastasis-associated protein 2 (MTA2), histone deacetylases 1 (HDAC1) and Retinoblastoma binding protein 4 (RBBP4) are components of the nucleosome remodeling and deacetylase (NuRD) complex. The NuRD complex is responsible for transcriptional repression through histone deacetylation and nucleosome remodeling (25). Coincidently, NuRD complex core members, as the helicase CHD4, was also found to be an interactor and epigenetic coregulator of PAX3-FOXO1 in alveolar rhabdomyosarcoma (26).

To prioritize the putative PAX8-interacting partners for further study, we ranked the peptides by confident identification score (i.e. MaxQuant score), correlation with PAX8 expression in ovarian tumor tissues, and co-dependency in ovarian carcinoma cell lines. SOX17, a member of the Sry-related HMG box transcription factors family, exhibited the strongest correlation, co-dependency with PAX8 in ovarian cancer, and was identified among the top-ranked most abundant putative PAX8-interacting partners (Table 1 and SI 1). We confirmed SOX17 as a *bona fide* PAX8-interacting partner by showing co-immunoprecipitation using agarose beads covalently bound to either a specific anti-PAX8 or anti-SOX17 antibody, compared to rabbit IgG control beads (Figure 1E), indicating that PAX8 and SOX17 physically interact. Consistent with this finding, we observed co-elution of PAX8 and SOX17 in the same large molecular size fractions (Figure 1F), indicating that they are part of the same complex. We did not observe monomeric PAX8 or SOX17 in lower molecular weight fractions.

### PAX8 physically interacts with SOX17 in high-grade serous ovarian carcinoma

We next characterized the location and levels of expression of PAX8 and SOX17 in five normal human fallopian tube tissues compared to five different HGSOC cases by immunohistochemistry. As shown in Figure 2A, we observed the co-expression of PAX8 and SOX17 restricted to the FT secretory epithelial cells. We also observed the over-expression of both PAX8 and SOX17 in all HGSOC cases (Figure 2A). Interestingly, PAX8 and SOX17 gene expression levels are significantly higher in ovarian cancers and in benign fallopian tubes than normal ovaries (SI 2A-2B-2C).

**Figure 2:**
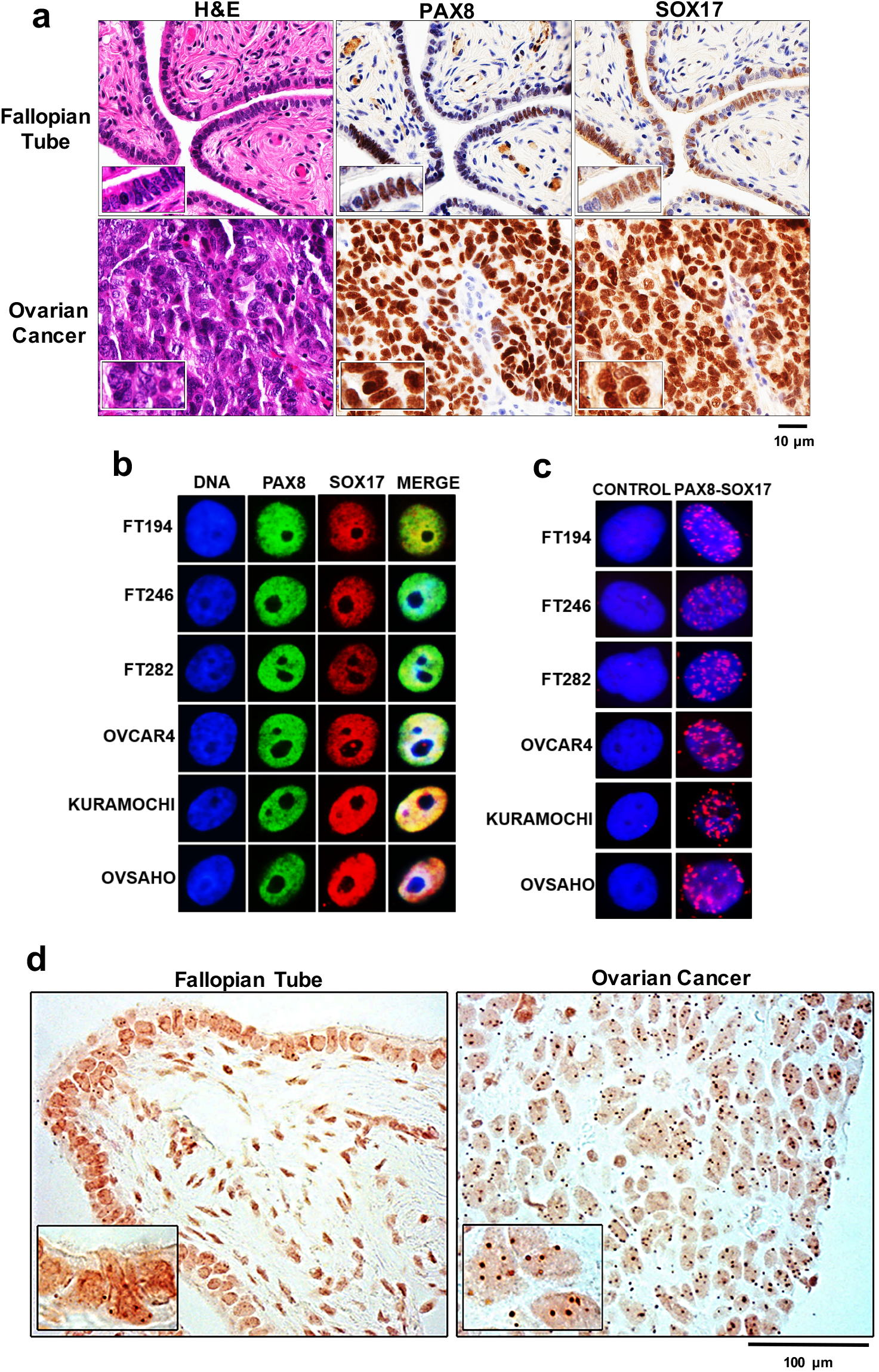
PAX8 physically interacts with SOX17 in the fallopian tubes and in ovarian cancer. (a) Immunohistochemistry showing nuclear co-expression of PAX8 and SOX17 in FTSEC and HGSOC cells. 5 normal samples and 5 ovarian cancer patients were analyzed and one representative case of each is shown. (b) Immunofluorescence showing co-localization of PAX8 and SOX17 in benign cells (FT194, FT246, and FT282) and in malignant cells (OVCAR4, KURAMOCHI, and OVSAHO). (c) Proximity ligation assay signals in secretory cells (FT194, FT246, and FT282) and in carcinoma cells (OVCAR4, KURAMOCHI, and OVSAHO) are shown in red. The nuclei are stained with DAPI (blue). The nuclei were acquired in one z-plane with 60X magnification. (d) *In situ* proximity ligation assay signals in normal fallopian tube sections and high-grade serous ovarian cancer samples shown in brown and the nuclei in orange. The nuclei were acquired in one z-plane with 40x magnification. *Number of PLA signals (PAX8-SOX17 interactions) was determined by counting 100 nuclei per sample (Supplemental information 2).

Our immunohistochemical findings are supported by high-resolution immunofluorescence analyses showing the nuclear co-localization of PAX8 and SOX17 in three different immortalized fallopian tube secretory cell lines (FT194, FT246, and FT282) and three high-grade serous ovarian carcinoma cell lines (OVCAR4, KURAMOCHI, and OVSAHO) (Figure 2B). To confirm the PAX8-SOX17 physical interaction, we performed an *in situ* proximity ligation assay (PLA), which allows the identification of both stable and transient interactions at native protein levels. We confirmed increased interaction between PAX8 and SOX17 in all tested high-grade serous cells: OVCAR4, KURAMOCHI, and OVSAHO (Figure 2C and SI 2D). As expected for transcription factors, the observed protein-protein interactions were nuclei localized. Moreover, we further explored the PAX8-SOX17 interaction in five different HGSOC tissue samples and again we observed a stronger and higher number of PLA signals in the cancer samples than in the normal fallopian tube samples (Figure 2D), confirming that the PAX8-SOX17 interaction may be important in the process of malignant transformation. Moreover, in the normal fallopian tube cases, the PLA signals (PAX8-SOX17 interaction) were restricted to the secretory cells, reinforcing the hypothesis that these cells are the site of origin for HGSOC.

### PAX8 and SOX17 are transcription co-regulators

To assess whether SOX17 and PAX8 could mutually regulate each other, we used RNA interference to knockdown each gene individually. After PAX8 knockdown, SOX17 protein levels were significantly reduced, in all tested FTSEC and HGSOC cell lines, and actually became undetectable in some lines (Figure 3A-3B and SI 3). Conversely, SOX17 knockdown also led to a decrease in the PAX8 protein level, but the effects were much less pronounced. At the RNA level, PAX8 loss led to a significant decline of SOX17 expression in all cell lines studied (Figure 3C) and SOX17 knockdown similarly led to a decrease of PAX8 expression (Figure 3D). These data suggest that SOX17 and PAX8 can transcriptionally regulate each other’s mRNA expression and that, in the absence of PAX8, SOX17 protein is rapidly depleted.

**Figure 3:**
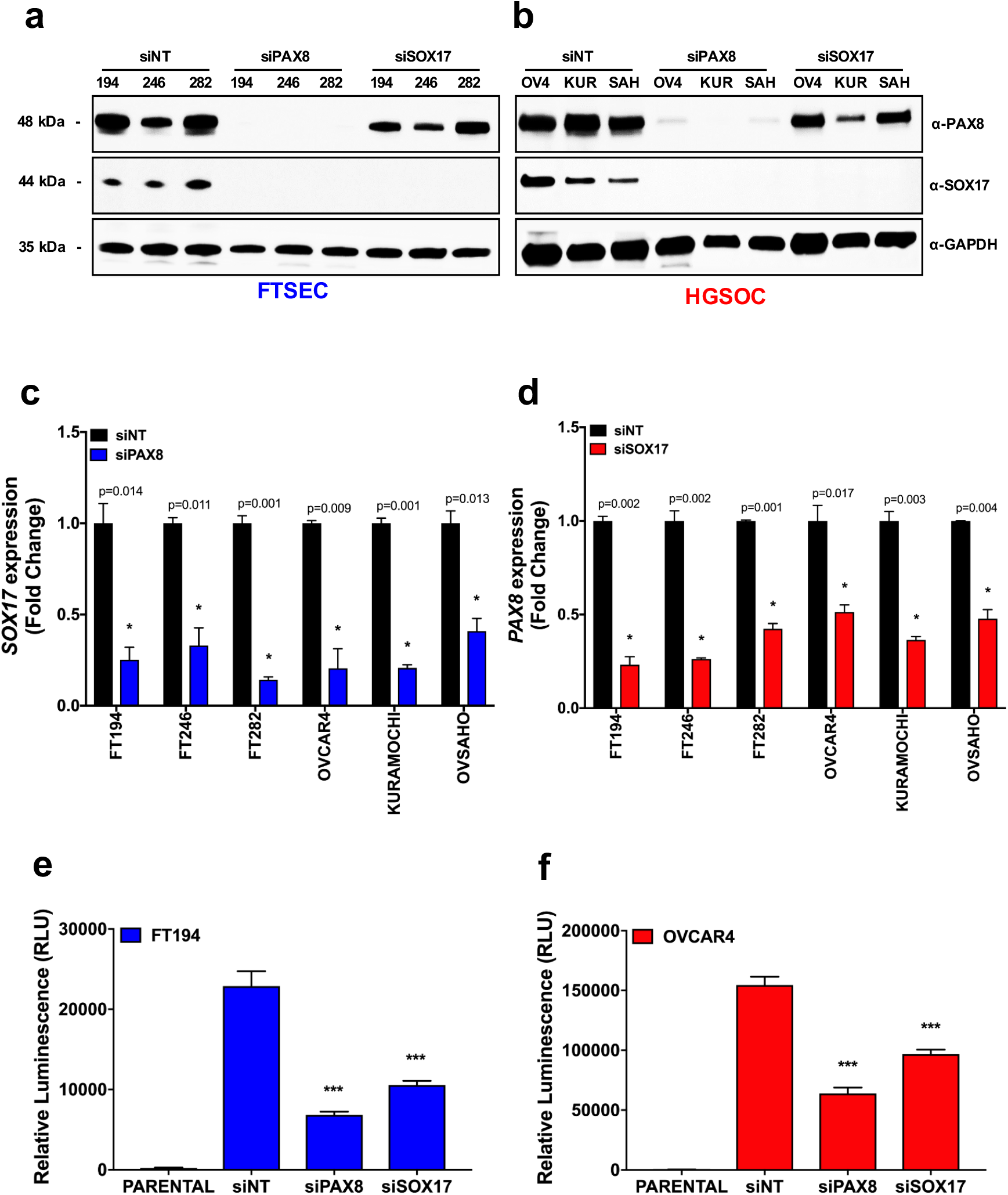
PAX8 and SOX17 are transcription co-regulators. (a-b) Immunoblot analyses following knockdown of PAX8 or SOX17 in FTSEC (FT194, FT246, and FT282) and in HGSOC (OVCAR4, KURAMOCHI, and OVSAHO) cells showing SOX17 expression dependency on PAX8 regulation. (c-d) Real-time PCR analysis following of PAX8 or SOX17 in FTSEC (FT194, FT246, and FT282) and in HGSOC (OVCAR4, KURAMOCHI, and OVSAHO) cells depicting the transcriptional co-regulation of SOX17 by PAX8. (e-f) Luciferase reporter assay using reporter with 5X PAX8-recognition sequence.

Since the PAX8-SOX17 complex appears to transcriptionally regulate the PAX8 and SOX17 promoters, we performed luciferase reporter assay to directly test the transcriptional effect of this complex on minimal promoters containing five times PAX8 binding sites (27). Consistent with the results described above, knockdown of either PAX8 or SOX17 demonstrated a significant decline of luciferase activity mediated by PAX8-binding sites (Figure 3E-3F).

### PAX8 and SOX17 regulate a common set of genes

To determine which pathways are regulated by the PAX8-SOX17 complex, we first performed RNA-seq following depletion of each factor individually, and in combination (siDUAL). Control cells received a non-targeting siRNA control pool (siNT). The efficiency of the knockdowns was assessed both by the number of sequenced reads, and Western blot (SI 3). Unsupervised analysis of the significantly altered transcripts after the loss of PAX8, SOX17, or both together is shown in Figure 4A and Table 2. PAX8 and SOX17 can both negatively and positively regulate gene expression. PAX8 target genes were significantly more likely to be regulated by SOX17 than expected by chance (p<0.00001, Chi-squared test). Treatment with siRNAs to simultaneously deplete both factors largely phenocopied the maximal effect of either siPAX8 or siSOX17 (Figure 4A). We focused on the 380 genes that were commonly up-regulated (Figure 4B) under all three conditions. These genes were enriched in pathways associated with cell adhesion, blood vessel development, and angiogenesis (Figure 4C).

**Table 2:** Top PAX8-SOX17 commonly regulated genes.

| COMMONLY UP-REGULATED TARGET GENES |  | COMMONLY DOWN-REGULATED TARGET GENES |  |
| --- | --- | --- | --- |
| Gene | Log-fold change | Gene | Log-fold change |
| SERPINE1 | 5.54863858 | PYY | -8.0262675 |
| IL11 | 5.27536243 | TMEM101 | -5.7263682 |
| NPPB | 5.18800181 | FGF18 | -5.4123449 |
| KLK5 | 4.50271261 | TGM7 | -4.9926057 |
| FGF1 | 4.4055198 | CA2 | -4.8188457 |
| CPA4 | 4.32537639 | CCNA1 | -4.5055642 |
| MYL7 | 3.80515773 | CXCL5 | -4.409147 |
| ADAMTSL4 | 3.79017684 | FOXQ1 | -4.3460707 |
| FILIP1L | 3.64159103 | NOTUM | -4.3070766 |
| HBEGF | 3.62339664 | GBP5 | -4.2098481 |
| TGFBI | 3.61350996 | TMEM171 | -4.1579087 |
| RASD2 | 3.52832382 | ZBED6CL | -4.0422849 |
| PMEPA1 | 3.48254273 | BMPER | -3.7191215 |
| SEMA3C | 3.43402726 | FST | -3.4878197 |
| DYSF | 3.32114739 | ATP2B2 | -3.4437388 |
| ITGA2 | 3.314082 | LRRC61 | -3.441764 |
| CYP1B1 | 3.28587766 | RHOC | -3.4246182 |
| NCF2 | 3.26470577 | MYB | -3.4111303 |
| ABCC3 | 3.21212074 | NMU | -3.4026155 |
| DAB2 | 3.18763588 | PRRX2 | -3.3958186 |
| CA9 | 3.17582117 | TXK | -3.3917272 |
| CD274 | 3.17245293 | AGMAT | -3.3704743 |
| CSF1R | 3.16372011 | CTTNBP2 | -3.3622138 |
| AKR1B10 | 3.09123637 | IQGAP2 | -3.300818 |
| SULT2B1 | 3.0772257 | KLHL14 | -3.2936874 |
| PDCD1LG2 | 3.06482259 | LRRC41 | -3.290006 |
| PCP4L1 | 3.05895596 | IMPDH1 | -3.285422 |

**Figure 4:**
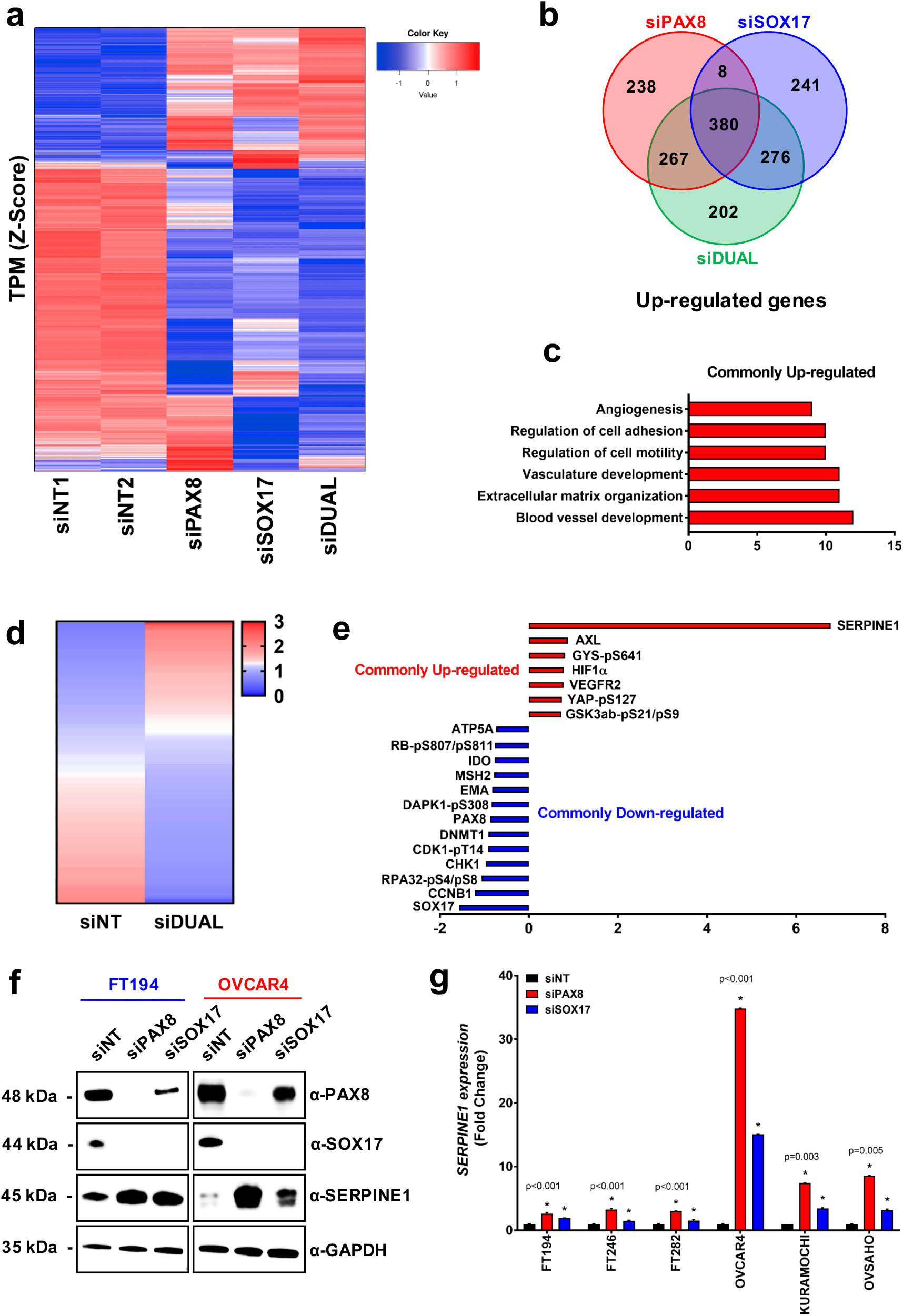
PAX8 and SOX17 regulate a common set of genes. (a) RNA-seq unsupervised clustering of PAX8, SOX17, and commonly regulated target genes. Profiles obtained with OVCAR4 after 72 hr knockdowns; fold change >1; *P* < 0.05. (b) Venn diagram representing number of genes up-regulated under each condition. (c) Ontology analysis of the PAX8-SOX17 commonly up-regulated genes. (d) RPPA unsupervised clustering of PAX8-SOX17 commonly regulated proteins. (e) Top-ranked PAX8-SOX17 commonly regulated proteins. (f) Immunoblot showing SERPINE1 up-regulation after PAX8 or SOX17 knockdown. (g) Real-time PCR showing *SERPINE1* up-regulation after PAX8 or SOX17 knockdown.

In order to further characterize the gene regulation coordinated by PAX8/SOX17, we performed targeted functional proteomic profiling using reverse-phase protein arrays (RPPA)s. Knockdown of PAX8, SOX17 or both resulted in significant changes across 142 proteins (Figure 4D and Table 3). Consistent with our observations made using RNA-seq, ontology analysis of the identified target genes showed that cell adhesion, and angiogenesis were among the pathways most significantly altered after PAX8 and/or SOX17 loss (SI 4C). When examining individual genes, we found that Serpin Family E Member 1 (SERPINE1), also called Plasminogen Activator Inhibitor 1 (PAI1), was the most highly elevated protein following knockdowns (Figure 4E). The protein data corroborate the RNA-seq analysis which also found SERPINE1 mRNA levels the most commonly up-regulated (SI 4A-4B). Interestingly, this gene has been implicated as an inhibitor of the tissue-type plasminogen activator and angiogenesis (28). Using immunoblotting and quantitative-PCR, we confirmed that PAX8 or SOX17 knockdown significantly increased the expression of SERPINE1 and all tested cells (Figure 4F-4G). Moreover, SERPINE1 presented a strong differential expression pattern between benign and malignant cells. All FTSEC lines exhibited higher levels of SERPINE1 compared to isogenic oncogene-transformed lines or ovarian cancer cell lines (SI 4D) and this expression was inversely correlated with the levels of PAX8/SOX17 (SI 4E). This suggests an important role for PAX8-SOX17-mediated SERPINE1 suppression in malignant transformation.

**Table 3:**
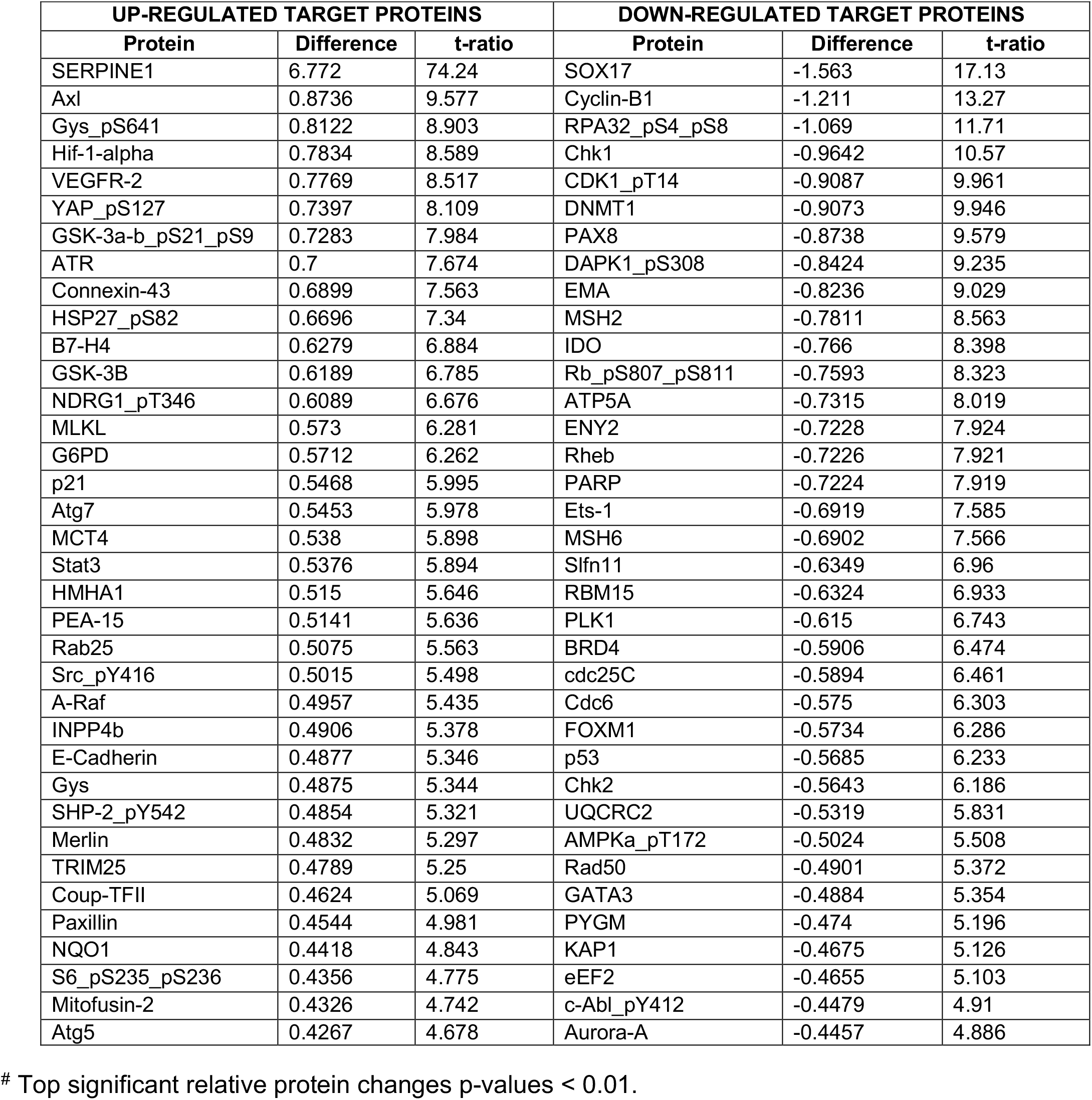
PAX8-SOX17 significantly co-regulated target proteins

### SERPINE1 participates in PAX8-SOX17-mediated angiogenesis

Using an angiogenesis antibody array, we further examined the levels of 35 different secreted angiogenesis mediators in a panel of human ovarian carcinoma cells (OVCAR3, OVCAR4, KURAMOCHI, and OVTOKO) and a panel of human fallopian tube secretory cells (FT33, FT194, FT246, and FT282). The FTSEC line-conditioned media exhibited a higher concentration of the angiogenesis inhibitors such as SERPINE1 and THBS1, while the ovarian cancer lines secreted more angiogenesis inducers such as VEGFA, CXCL8, and CXCL16 (Figure 5A and SI 5). We found that some angiogenic factors were regulated by PAX8 and SOX17. Indeed, VEGFA, CXCL8, and CXCL16 exhibited reduction after the knockdown of PAX8 or SOX17 in the cancer lines, while their secretion in the normal lines was not detected (Figure 5B). However, the most prominent effect observed in cancer cell lines was a large increase in SERPINE1 secretion following PAX8 knockdown. SERPINE1 is particularly interesting because its secretion levels were also elevated in FTSECs compared to ovarian cancer lines. Indeed, FTSEC-conditioned media presented an average of 20 ng/mL of SERPINE1, which was many times higher than the secretion from ovarian cancer lines, an average of 0.2 ng/mL (Figure 5C-5D). Corroborating our previous findings, the secretion of SERPINE1 was significantly increased in HGSOC conditioned media following PAX8 or SOX17 knockdown (Figure 5E-5F).

**Figure 5:**
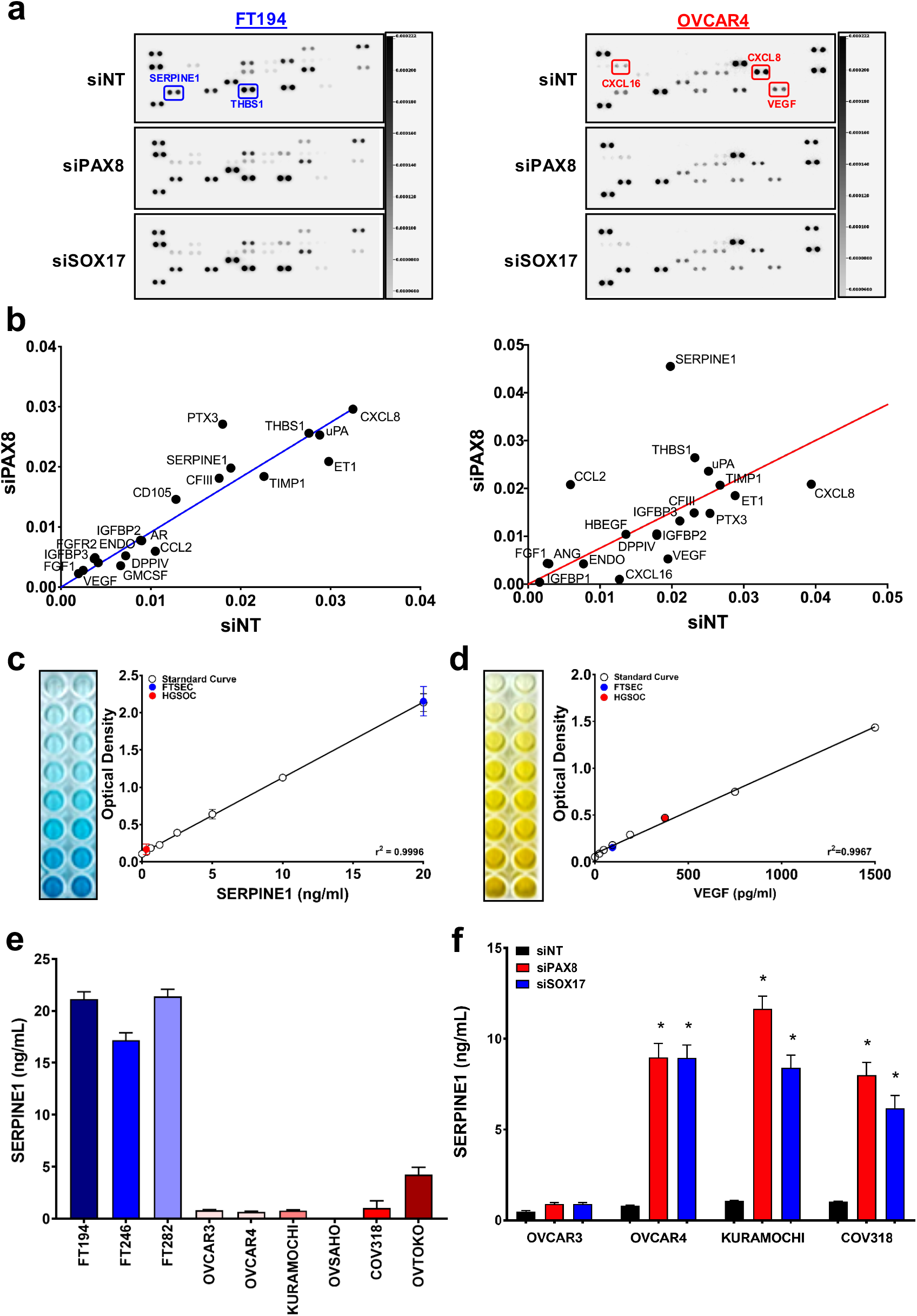
PAX8 and SOX17 regulate the secretion of angiogenesis mediators. (a) Human angiogenesis array of conditioned media from FTSEC and HGSOC cells following PAX8 or SOX17 knockdown. (b) Effect of PAX8 knockdown on specific analytes was produced by quantifying the array membrane spots intensity. (c) ELISA for quantification of secreted SERPINE1 in the FTSEC- and HGSOC-conditioned media. (d) ELISA for the quantitation of secreted VEGF in the FTSEC and HGSOC conditioned media. (e) ELISA showing SERPINE1 secreted by FTSEC and HGSOC. (f) ELISA experiments showing the effects of PAX8 or SOX17 knockdown on levels of SERPINE1 in HGSOC.

### Roles of SERPINE1 and its regulators, PAX8 and SOX17, on ovarian tumor angiogenesis

Using the human umbilical vein endothelial cells (HUVEC) tube-formation assay, we found that conditioned media from ovarian carcinoma cells induced the formation of blood vessels *in vitro*, although not to the same extent as recombinant VEGF. This effect was almost abolished in conditioned media from ovarian cancer lines with PAX8 or SOX17 knockdown (Figure 6A). However, no induction of HUVECs tube formation was observed with the FTSEC lines conditioned media, which exhibit a higher concentration of SERPINE1 (Figure 6A-6B). We further investigated the VEGFR2 pathway status and corroborated the inactivation of the downstream effectors PLCy1 and ERK1/2 in endothelial cells exposed to HGSOC conditioned media after the loss of PAX8 or SOX17 when compared with the no-targeting control knockdown (SI 6).

**Figure 6:**
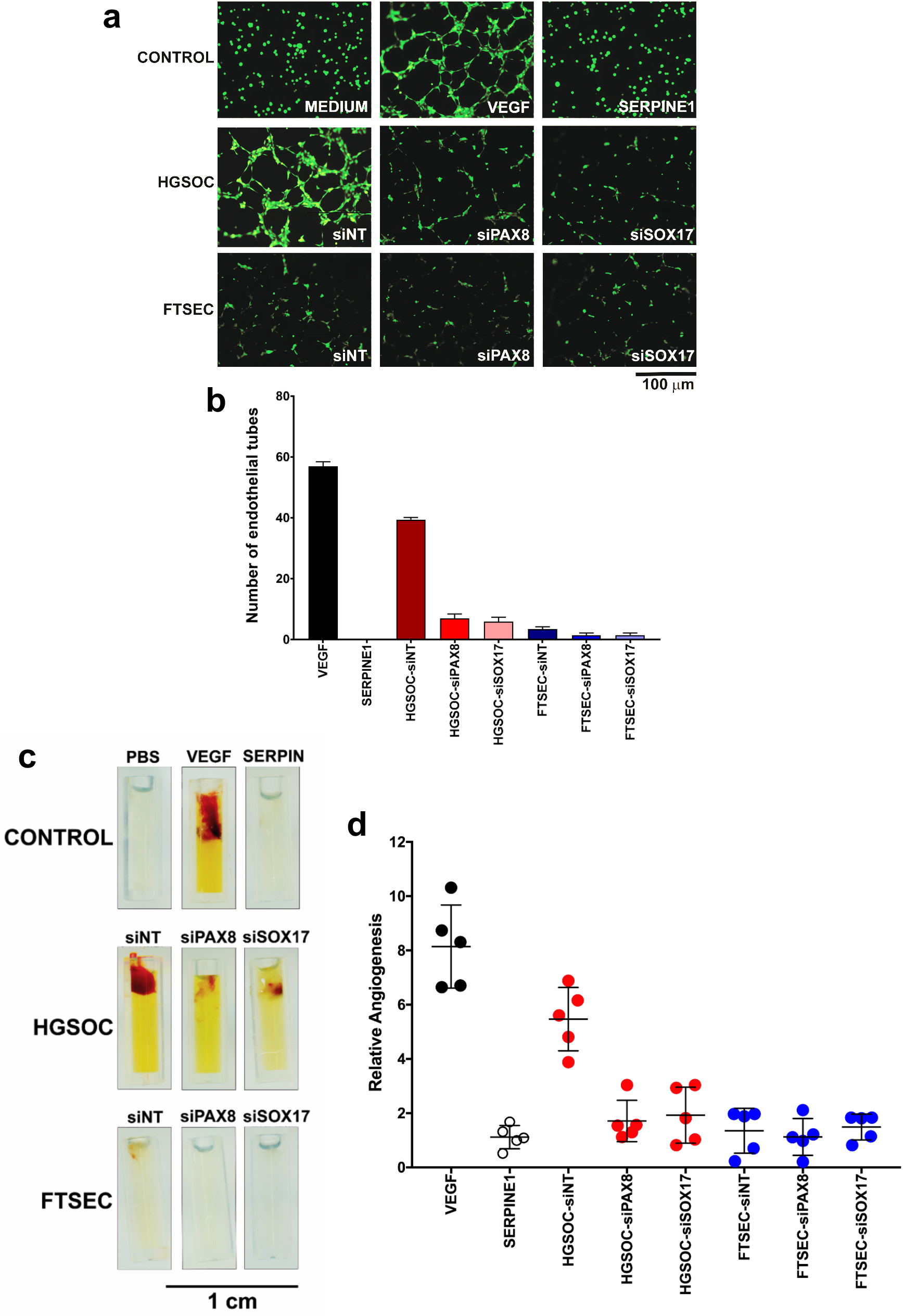
PAX8 and SOX17 promote ovarian cancer angiogenesis. (a) Endothelial cells tube formation assay. Representative images depict negative control, VEGF, SERPINE1, FTSEC, and HGSOC conditioned media after PAX8 or SOX17 knockdown. (b) Quantitation of the HUVEC neo-vessels loops. (c) Neovascularization in angioreactors containing conditioned media from HGSOC cells, but not from FTSEC after implantation in nude mice. (d) Quantitation of endothelial cell invasion into angioreactors.

To extend these findings to an *in vivo* system, we performed in nude mice the directed *in vivo* angiogenesis assay (DIVAA) (29, 30). The DIVAA assay uses semi-closed small silicone cylinders known as angioreactors. The cylinders can be filled with angiogenic or antiangiogenic compounds of interest. Following subcutaneous implantation in nude mice, host vascular endothelial cells will migrate into the angioreactors and proliferate to form new blood vessels, if the compound of interest is angiogenic. In the presence of VEGF, used as a positive control, a strong induction of angiogenesis was observed (Figure 6C-6D). No significant blood vessels were observed in the vehicle (PBS or Fresh Media) control or with SERPINE1. Cylinders containing HGSOC cell lines conditioned media revealed extensive angiogenesis. The presence of erythrocytes inside the newly developed blood vessels indicated that they were functional. In contrast, conditioned media from HGSOC cell lines with PAX8 or SOX17 knocked down showed a significant decrease in ovarian cancer-induced neovascularization. These results show that ovarian cancer cell lines have the capacity to induce angiogenesis *in vivo*, and that PAX8 and SOX17 participate in this process probably through SERPINE1 down-regulation.

### Knockdown of PAX8 or SOX17 decreases tumor formation in an ovarian cancer xenograft model

Because we observed significant and reproducible effects of PAX8/SOX17 on angiogenesis, we hypothesized that knockdown of these genes may reduce the capacity of malignant cells to develop tumors *in vivo*. We employed a doxycycline-inducible lentiviral shRNA system in order to express shRNAs against PAX8 or SOX17 to deplete PAX8 or SOX17 levels in 3 different cancer cell lines (OVCAR3, OVCAR4, and OVTOKO). Unfortunately, OVCAR3 and OVCAR4 cells appeared to be exquisitely sensitive to the loss of PAX8 and SOX17 and we were unable to identify clones with acceptable levels of down-regulation. However, some OVTOKO clones exhibited efficient PAX8 or SOX17 knockdown and up-regulation of SERPINE1 after the induction with doxycycline (Figure 7A). OVTOKO-shRNA harboring cells were injected intraperitoneally into NSG mice and incubated for 72 hours before doxycycline was added to the diet. After two weeks, control mice exhibited a significant tumor burden (Figure 7B). Doxycycline treatment reduced the tumor burden further to an almost undetectable level with no signs of ascites (Figure 7C-7D-7E), reinforcing our hypothesis that PAX8/SOX17 are modulating tumor angiogenesis by suppressing anti-angiogenesis factors as SERPINE1 and THBS1 (SI 7). These data show that inhibition of PAX8/SOX17 profoundly impairs the growth of ovarian carcinoma cells in mice, part of these effects likely due to suppression of tumor angiogenesis.

**Figure 7:**
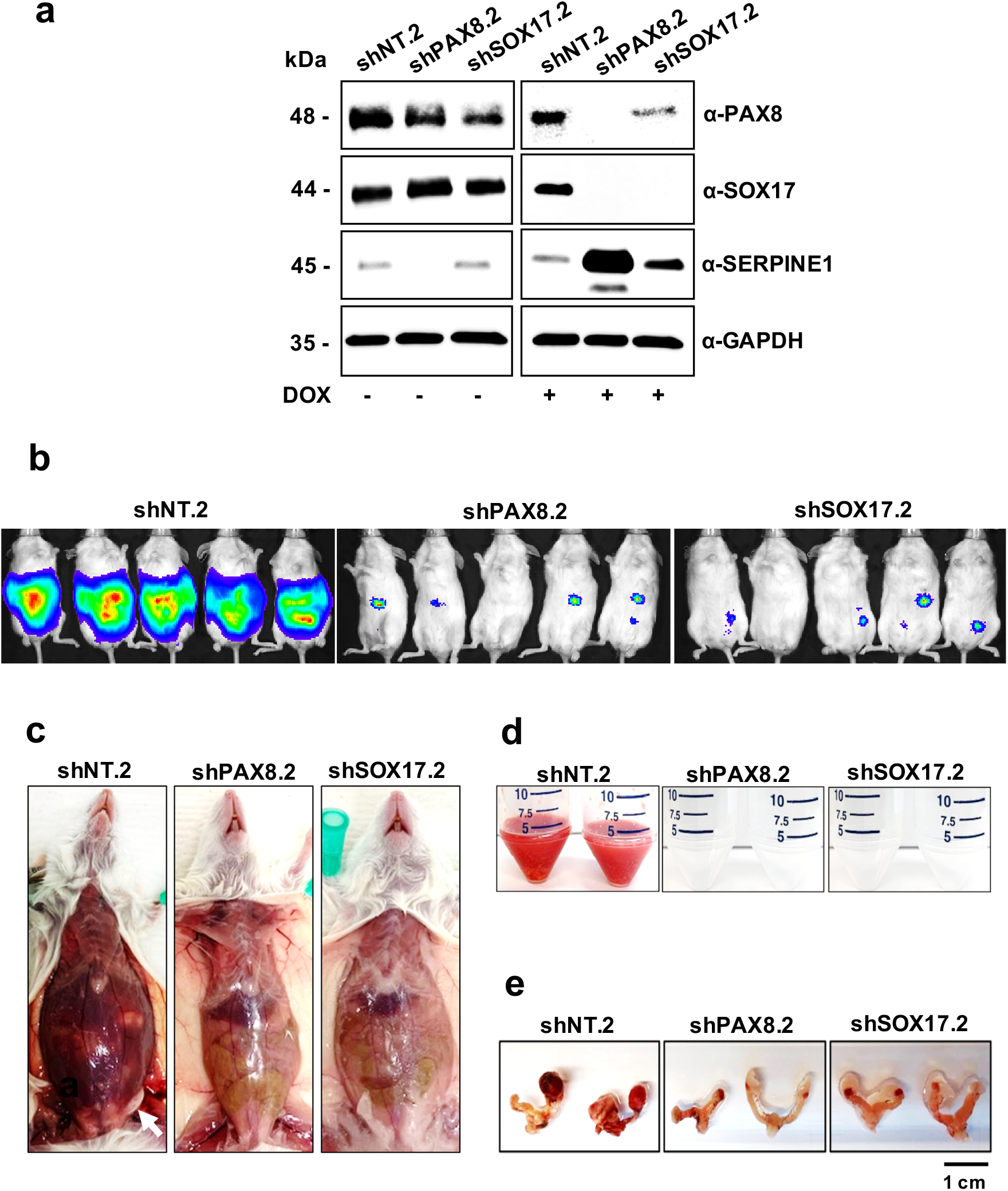
PAX8 or SOX17 doxycycline-inducible knockdown inhibits ovarian cancer progression. (a) Immunoblot of OVTOKO cells harboring inducible shPAX8 or shSOX17 before and after the induction with 1μg/ml of doxycycline. (b) *In vivo* imaging of OVTOKO tumors in NSG female mice. Animals were imaged after two weeks of doxycycline supplementation. (c) Necropsy of animals depicting ascites and tumors (white arrow) only in the non-targeting control group. (d) The volume of ascites collected from mice after two weeks of doxycycline supplementation. (e) NSG mice reproductive system depicting ovarian cancer volume.

## DISCUSSION

Many studies have implicated PAX8 as an important transcription factor in ovarian cancer. However, its mechanism of action in this regard remains unclear. Here, using a combination of biochemistry, *in vitro* models, and animal studies we make two important observations about how PAX8 governs tumor development. We show that PAX8 interacts with another developmental transcription factor, SOX17. PAX8 regulates expression of SOX17 and together they mediate a transcriptional program that drives angiogenesis in the malignant setting. Depletion of these proteins greatly attenuates angiogenesis in multiple systems.

Previous studies have shown that benign and malignant cells are distinguished by marked remodeling of the PAX8 cistrome and this implies that PAX8 may acquire new targets or functions in the malignant state (22). To investigate if the PAX8 re-distribution in cancer cells was due to changes in the PAX8 network, and to further clarify its roles in ovarian cancer, and possibly identify novel therapeutic opportunities, we first sought to identify its interacting partners in benign and malignant cells. Interestingly, its crucial role in transcriptional regulation was highlighted by our finding that multiple chromatin remodeling proteins interact with PAX8. Mechanisms of gene regulation by PAX8 are clearly complex as we found that, following PAX8 knockdown, a vast number of genes were up-regulated or down-regulated in these cells. A large number of the nucleosome remodeling molecules identified here are subunits of the NuRD complex, such as CHD4, MTA2, GATAD2A, GATAD2B, HDAC1, and RBBP4 suggesting that PAX8 participate in this chromatin-remodeling complex. Reinforcing these findings, a recent drug screen showed that PAX8 levels were strongly decreased by inhibitors of histone deacetylase (HDAC1) (31). Among the interacting partners identified, SOX17 was one of the mostly highly enriched in our mass spectrometry data. In addition, it exhibited a strong correlation and co-dependency with PAX8 in ovarian cancer using TCGA gene expression data. SOX17 is a transcription factor and member of the SOXF family, which has an HMG, β-catenin binding and transactivation domains (32). SOX17 biological function is always dependent on a dimerization partner (33) that is dynamic and specific to the cell context. SOX17-interacting partners can engage differentially in the genome, regulating different sets of genes (34). Therefore, we hypothesized that the PAX8-SOX17 transcriptional activity may be crucial in ovarian tumorigenesis. We found that both PAX8 and SOX17 exhibit expression in the nuclei of both benign and malignant secretory cells, where they physically and functionally interact. Our findings are in agreement with the current literature showing that PAX and SOX members can engage in transcriptional regulation. Indeed, PAX3 and SOX10 can physically interact and synergistically regulate MITF and c-RET enhancers (35). The PAX3-SOX10 interaction is important for melanoma cells, where these factors regulate cell motility, apoptosis, and proliferation (36). Additionally, PAX6 and SOX2 are also interacting partners in early neural differentiation and are necessary for neural progenitor cell pluripotency (37). Furthermore, PAX6 and SOX2 act as an oncogene and can induce cancer cell stemness (38).

We found that PAX8 and SOX17 can mutually regulate each other at the transcriptional level. At the protein level, PAX8 knockdown led to an almost complete disappearance of SOX17, and SOX17 knockdown led to a significant decrease in PAX8 levels. These results are reinforced by our recent findings showing that PAX8 and SOX17 were found to be master transcription factors that occupy regulatory elements related to their own encoding genes in ovarian cancer (39). Globally, PAX8 and SOX17 genomic binding sites co-localize within candidate active enhancers in HGSOC cell lines. In addition, PAX8 binds with high intensity near SOX17 gene locus, which confirms the co-regulation observed in SOX17 transcript and protein levels (39). Here, using transcriptomic and proteomic analyses we showed that PAX8 and SOX17 commonly regulate a family of genes associated with blood vessel formation, suggesting a cooperative role in orchestrating an important pro-angiogenic transcriptional program in ovarian cancer. In this setting, it is interesting to note that murine Sox17 can promote tumor angiogenesis and is regulated by the Notch pathway, known to contribute to angiogenesis (40–43).

Angiogenesis is a dynamic process rigorously regulated during embryogenesis, tissue regeneration, and ovulation. However, angiogenesis can also abnormally occur during malignant transformation (44, 45). Most of the angiogenesis mediators are secreted cytokines, but matrix metalloproteinases and integrins also participate in new blood vessel development (46). The most commonly secreted mediator in tumor angiogenesis is Vascular Endothelial Growth Factor (VEGF). VEGF acts through its receptor (VEGFR2) and downstream effectors such as PLCy1 and ERK1/2. Furthermore, VEGF signaling pathway abrogation has emerged as an attractive strategy of new targeted anti-cancer therapy (47). Indeed, we found that PAX8 and SOX17 loss triggered a reduction in angiogenesis both *in vitro* and *in vivo* model. We also observed a drastic decrease in tumor xenograft growth following PAX8/SOX17 knockdown, and while we cannot rule out the roles of other cell intrinsic processes, the decrease in angiogenesis is likely an important factor. It has been reported that patients with high levels of VEGF have the worst prognostic and lower survival rates. Moreover, ovarian cancer cells express on their surface VEGF receptors that contribute to angiogenesis and malignant progression (48). The VEGF pathway is also responsible for the formation of ascites in patients with ovarian tumors due to the elevated vascular permeability within peritoneal lining in the abdominal cavity (49). Consistent with this fact, we found a significant decrease in ascites formation following knockdown of PAX8 and SOX17.

In trying to better understand the mechanisms of angiogenesis orchestrated by PAX8 and SOX17, we examined individual genes and identified those that were most affected by the knockdowns. We found SERPINE1 to be the most highly commonly up-regulated gene both at the transcript level (using RNA-seq) and protein level (through RPPA). SERPINE1 is a serine proteinase inhibitor, belonging to the Serpin family, which is an important endothelial plasminogen activator inhibitor and urokinase inhibitor (50). Important roles in coagulation, extracellular matrix remodeling, and angiogenesis have been reported for SERPINE1. The anti-angiogenic effects of SERPINE1 seem to be mediated by binding to vitronectin and blocking αvβ3 and uPAR binding sites (51). Therefore, binding of secreted SERPINE1 to vitronectin blocks cell adhesion, migration, and inhibits angiogenesis (52). Moreover, vitronectin and integrin-αvβ3 strongly intensify VEGFR2 activation by VEGF (53). SERPINE1 acts as an angiogenesis inhibitor by blocking the vitronectin-integrin-VEGFR2 axis and simultaneous abrogation of the VEGF-signaling pathway. Our set of experiments involving endothelial cell tube formation *in vitro* and the migration of endothelial cells into subcutaneous implants *in vivo* suggest that SERPINE1 is an essential inhibitor of tumor angiogenesis and is under PAX8-SOX17 regulation. Based on these experiments, we hypothesize that suppressing SERPINE1 through malignant transformation, triggers VEGFR2 activation *in vivo* and contributes to tumor angiogenesis (SI 7).

Our results may also help explain some of the developmental defects described in the *Pax8* knockout mouse (14). *Pax8^−/−^* mice are infertile because they lack a functional uterus revealing only remnants of myometrial tissue. In addition, the vaginal opening is absent. Folliculogenesis, ovarian hormone production, and transcription of pituitary hormones are in a normal range. Thus, infertility in *Pax8^−/−^* mice seems to be due to a defect in development of the reproductive tract rather than to hormonal imbalance, pointing to a direct morphogenic role for Pax8 in uterine development. Our observation that Pax8 and Sox17 orchestrate an angiogenic program may help explain the atresia of the reproductive tract seen in the *Pax8^−/−^* mice. The absence of Pax8 in the developing reproductive tract is likely accompanied by low Sox17 and high Serpine1. These conditions would effectively shut down blood vessel development and prevent the development of the organ. This is reminiscent of the severe embryopathy seen with thalidomide in the early 1960s (54). Thalidomide was marketed as an anti-emetic which was later shown to have anti-angiogenic properties that cause severe birth defects, including phocomelia (limb defects), genital, and internal organ absence or malformation.

In summary, we have shown for the first time that PAX8 physically interacts with SOX17 in FTSEC and HGSOC leading to changes in multiple transcriptional programs, including modulation of genes mediating tumor angiogenesis. Further analyses of the genes regulated by PAX8/SOX17 identified SERPINE1 as an anti-angiogenic factor that is suppressed by PAX8-SOX17 in cancer cells. Knockdown of PAX8/SOX17 in this setting results in a decrease of blood vessel formation. Using cell-based and animal models, we then functionally demonstrate that PAX8/SOX17 can regulate angiogenesis during tumor development. Inhibition of tumor angiogenesis, perhaps through PAX8/SOX17 pathway inhibition, has potential value as part of an anti-angiogenic approach to the treatment of ovarian cancer (40, 55).

## METHODS

### Cells and Tissues

Human immortalized fallopian tube secretory cells (FT33, FT194, FT246, and FT282) were maintained in standard conditions as previously described (56) and grown in DMEM/F12 containing 2% Ultroser-G serum substitute. Human ovarian cancer cells were acquired from different sources. OVCAR3 was obtained from ATCC and maintained as recommended. OVCAR4 was acquired from William C. Hahn’s lab-CCLE (Dana-Farber Cancer Institute) and grown in RPMI 1640 containing 10% fetal bovine serum. KURAMOCHI, OVSAHO, and OVTOKO were obtained from JCRB and maintained as recommended. The human umbilical vein endothelial cells (HUVEC) were acquired from Sigma-Aldrich and maintained according to manufacturer instructions. All cell lines were sent to the Wistar Institute Genomic Core Facility for authentication using short tandem repeat profiling and for detection of *Corynebacterium bovis* infection. In addition, all cell lines were also tested for *Mycoplasma* sp. at the University of Pennsylvania Cell Center.

Following approval by the Hospital of Pennsylvania Institutional Review Board, we obtained human fallopian tube and human high-grade serous ovarian carcinoma formalin-fixed and paraffin-embedded sections from the Department of Pathology at HUP to evaluate the expression of PAX8 and SOX17.

### Purification of endogenous PAX8

Benign and malignant cells were grown in 15-cm plates until 90% confluence, washed twice with PBS, trypsinized, neutralized and collected. Nuclear fractionation was prepared as previously published (24). Harvested cells were resuspended in hypotonic buffer [20 mM HEPES, pH 7.5, 10 mM KCl, 1 mM EDTA, 1 mM EGTA, 1.5 mM MgCl_2_, 2 mM dithiothreitol, protease inhibitors (Sigma-Aldrich: P8340), and phosphatase inhibitors (Sigma-Aldrich: P5726)] and incubated for 30 minutes. Samples were then disrupted through a 22G needle and centrifuged at 10,000 × *g* for 10 minutes at 4°C. Nuclei-enriched fraction was sonicated with complete RIPA buffer (Cell Signaling: 9806S) containing protease inhibitors (Sigma-Aldrich: P8340), and phosphatase inhibitors (Sigma-Aldrich: P5726), and spun down for 10 minutes at 10,000 × g at 4°C. The supernatant (500 μg of nuclear extract) was incubated for 16 hours at 4°C with 105 μg of PAX8-specific antibody (Proteintech: 10336-1-AP) coupled to 1 ml of Protein A agarose resin (Thermo-Fisher: 44893) or with 105 μg of normal rabbit IgG (Proteintech: 30000-0-A) coupled to 1 mL of Protein A agarose resin, as negative control. The columns were washed three times with 10 mL of 0.1 M phosphate and 0.15 M NaCl, pH 7.2 and eluted with 0.5 M NaCl and 0.1 M glycine, pH 2.8. Fractions had their pH equilibrated with 1M Tris, pH 9.5, separated by gel electrophoresis, Coomassie blue-stained and lanes were sent for mass spectrometry analysis.

The affinity column eluates containing PAX8 were also loaded in 100 ml of Sephacryl S-300 column (Sigma-Aldrich: S300HR-100ML) equilibrated with 50mM Tris, pH 7.5, 100 mM KCL, 0.5 M NaCl, 1% NP-40 and 1% glycerol. We collected one hundred and fifty 500 μL fractions and protein peaks were separated by gel electrophoresis, silver-stained (Thermo-Fisher: 24600), checked by WBs for the presence of PAX8, and positive-samples were also sent for mass spectrometry analysis.

### PAX8 interacting-partners identification

Coomassie blue-stained lanes containing PAX8 were analyzed by nanoLC-MS/MS setup as previously described (57). In summary, HPLC gradient was set between 0-30% of solvent A = 0.1% formic acid and solvent B = 95% acetonitrile, 0.1% formic acid for one hour followed by five minutes of 30% to 85% of solvent B and ten minutes of isocratic 85% solvent B. Flow rate of nLC was set to 300 nL/min and coupled to Orbitrap Fusion Tribrid mass spectrometer (Thermo Fisher, USA) with 2.5 kV spray voltage and 275 °C of capillary temperature. Full mass spectrometry was performed using a resolution of 120,000 and 27 of HCD.

DDA files were analyzed with MaxQuant (58) using a SwissProt human database. iBAQ quantification was used for enrichment analysis and data were log2 transformed and normalized by subtracting the average of all valid values for each sample. Statistics analysis was obtained applying a two-tails heteroscedastic t-test.

### Co-immunoprecipitation

500 μg of nuclear lysates were incubated with 25 μg of a specific PAX8-antibody (Novus: NBP1-32440), 25 μg of a specific SOX17-antibody (Abcam: ab224637), or 25 μg of a normal rabbit IgG (Cell Signaling: 2729S) covalently-coupled to activated agarose beads (Thermo-Fisher: 26148) as manufacturer’s recommended protocol.

### Conditioned medium

Secretory cells and carcinoma cells were growth in 60 mm dish at 37 °C and 5% CO^2^. Conditioned media were retrieved by spinning down at 2,000 × g for 5 minutes at 4 °C then supernatants were passed through 0.22 μm filter (Millipore: SLGP033RS).

### Immunohistochemistry

Fallopian tube and high-grade ovarian carcinoma sections were processed as previously reported (59). The immunohistochemical staining were performed using a dilution of 1:500 of anti-PAX8 antibody (Novus: NBP1-32440) or a dilution of 1:500 of anti-SOX17 antibody (Novus: NBP1-47996). Slides were scanned with Aperio CS2.

### Immunofluorescence

Ten thousand FTSEC and HGSOC cells were seeded on imaging plates (Eppendorf: 0030741030) and allowed to grow for 24 hours. Following cells were washed twice in PBS and fixation was performed for 15 minutes on Paraformaldehyde 4% (Thermo-Fisher: AAJ19943K2) at room temperature. After the incubation period, cells were washed twice in PBS and permeabilized with Triton X-100 0.25% (Boston BioProducts: P-924) for 15 minutes. Aldehydes residues were quenched with Glycine 100 mM (Sigma-Aldrich: 50046-50G) for 15 minutes. The unspecific sites were blocked with a solution of 1% bovine serum albumin, and 0.1% Tween 20 in PBS for 30 minutes. Samples were incubated for 16 hours with a dilution of 1:500 of anti-PAX8 (Novus: NBP1-32440) or a dilution of 1:500 of anti-SOX17 (Novus: NBP1-47996) at 4 °C. Cells were then washed three times of 5 minutes each with a solution of 1% bovine serum albumin, and 0.1% Tween 20 in PBS followed by incubation for one hour with anti-mouse-AlexaFluor488 or anti-rabbit-AlexaFluor594. Cells were washed three time in PBS, mounted in Fluoromount-G with DAPI, and Images were acquired at 60X magnification employing a Nikon Eclipse Ti inverted microscope.

### *in situ* Protein-protein interaction analysis

Prior to the protein-protein interaction staining by *in situ* proximity ligation assay (PLA), tissue sections and cell lines were processed as described in the IHC and IF sections. PLA signals were determined employing Duolink Probes Anti-Mouse MINUS (Sigma-Aldrich: DUO92004) and Anti-Rabbit PLUS (Sigma-Aldrich: DUO92002) as manufacturer’s recommended protocol following overnight incubation with anti-PAX8 mouse antibody 1:250 (Novus: NBP2-29903) and anti-SOX17 rabbit antibody 1:250 (Cell Signaling: 81778S) at 4 °C. Red fluorescent signals was obtained using detection reagent red (Sigma-Aldrich: DUO92008) and chromogen signals was acquired using detection reagent brightfield (Sigma-Aldrich: DUO92012).

### siRNA knockdown

Fallopian tube secretory and ovarian carcinoma cells knockdowns were performed by reverse transfection with a mixture containing of 30 nM of ON-TARGETplus Human PAX8 siRNA (Dharmacon:L-003778-00-0005), ON-TARGETplus Human SOX17 siRNA (Dharmacon: L-013028-01-0005) or non-targeting siRNA as negative control (Dharmacon: D-001810-10-05), 3 × 10^5^ cells, 10 μl of Lipofectamine RNAiMax (Thermo-Fisher: 13778075), and Opti-MEM reduced serum medium (Gibco: 31985088) as recommended by the manufacturer’s protocol.

### Western blot analysis

Samples were incubated with RIPA buffer (Cell Signaling: 9806S) for 30 minutes at 4°C followed by centrifugation at 10,000 × g for 5 minutes. Supernatants’ protein concentration was estimated by BCA method (Thermo-Fisher: 23227). Thirty micrograms of each sample were mix with sample buffer loaded and separated using Mini-PROTEAN TGX 4-15% polyacrylamide gels (BioRad: 4561083) and Tris/Glycine/SDS buffer (BioRad: 1610732). TURBO transfer system (BioRad: 1704156) was employed to move separated samples from gels to PVDF membranes. Primary antibodies: anti-PAX8 (Novus: NBP1-32440), anti-SOX17 (Abcam: ab224637) or anti-GAPDH (Cell Signaling: 5174) were diluted 1:1000 in 5% nonfat milk in TBS containing 0.1% Tween 20 and incubated with the membranes overnight at 4°C. After the period of incubation, membranes were washed three times with TBS containing 0.1% Tween 20 then incubated with anti-Rabbit IgG-HRP conjugated (Cell Signaling: 7074). Images were acquired by chemiluminescence using Clarity ECL (BioRad: 1705062).

### Real-time PCR

Samples total RNA was purified using the RNeasy Plus Mini Kit (Qiagen: 74134), quantified and used as a template for the synthesis of single-stranded cDNA employing the High Capacity cDNA Reverse Transcription Kit (Thermo-Fisher: 4374966). To access the gene expression changes we employed the TaqMan Assay (Thermo-Fisher: 4331182) using 100 ng of cDNA per 20 μL of final reaction with TaqMan Fast Advanced Master Mix (Thermo-Fisher: 4444557) as recommended by manufacturer’s protocol.

### RNA-seq

Transcriptome analysis of OVCAR4 cells after PAX8, SOX17 or both knockdown simultaneously was performed as previously described (39, 60). In summary, ovarian carcinoma cells had their RNA chemically purified using the Nucleospin RNA Plus kit (Macherey-Nagel: 740984.50) as recommended by the manufacturer’s protocol. Poly-A non-stranded library were prepared using the newly extracted RNA and forty million reads were sequenced by BGI platform. Bioinformatic analyses were executed employing the R package DESeq2 (version 1.24.0). Significant changes were designated as log2 fold change ≥ 1 and adjusted p-value ≤ 0.01. Metascape tool was employed to identify the differentially enriched pathways.

### Reverse-phase protein array

Arrays were performed at the Department of Bioinformatics and Computational Biology in MD Anderson Cancer Center as previously described (61, 62). The platforms contains over 300 antibodies exclusively validated with a Pearson coefficient > 0.7 of correlation between RPPA and WB were employed in the proteomic analysis (63). Spots intensities were generated by colorimetric reaction employing the Dako Cytomation-Catalyzed System.

### Luciferase reporter assay

Briefly, half-million cells were co-transfected with 1 μg of PAX8-(Firefly) Luciferase reporter vector (27), 0.5 μg of CMV-(Renilla) Luciferase control vector (Promega: E2261) and 30 nM of PAX8, SOX17 or Non-targeting siRNA, using Lipofectamine 2000 (Thermo-Fisher: 11668027) as recommended by manufacturer’s protocol. Plates containing the different transfected cells were incubated for 24 hours at 37°C before the luciferase activity was measured using the Dual-Glo luciferase detection kit (Promega: E2920).

### Array and ELISA

Angiogenesis secreted mediators were identified from fallopian tube secretory cells and ovarian carcinoma cells conditioned media employing the Human Angiogenesis Antibody Array (R&D systems: ARY007). ELISA was employed for the precise quantification of VEGF (R&D systems: DVE00) and SERPINE1 (R&D systems: DSE100) also from cells conditioned media as recommended by the manufacturer’s protocol. Fresh DMEM/F12 or RMPI media were tested and considered as the negative control.

### *In vitro* angiogenesis assay

Twenty thousand HUVEC cells were seeded in reduced growth factor basement membrane extract (BME)-coated 96-well plate (R&D systems: 3470-096-K). Endothelial cells were exposed to 100 μl of the different benign or malignant cells conditioned media, 10 ng/ml VEGF (R&D systems: 293-VE-010) or 10 μg/ml SERPINE1 (R&D systems: 1786-PI-010) for 6 h at 37 °C. HUVEC cells were labeled with 2 μM Calcein AM as recommended by the manufacturer’s protocol to facilitate the image acquisition using a Nikon Eclipse Ti inverted microscope. The number of complete endothelial loops per field were counted and compared.

### *In vivo* angiogenesis assay

Following IACUC review and approval (Protocol #806687), *in vivo* angiogenic responses were analyzed by directed *in vivo* angiogenesis assay (DIVAA) (R&D systems: 3450-048-K) as recommended by the manufacturer’s protocol. Briefly, six-week female nude mice (JAX: 002019) were kept in aseptic condition under the Stem Cell and Xenograft Core barrier at the University of Pennsylvania. Mice cohorts were anesthetized with 2% isoflurane prior the subcutaneously implantation of angioreactors i.e. one-centimeter flexible silicone cylinder. Dorsal-lateral incisions were made on each nude mouse, where angioreactors filled with FTSEC or HGSOC conditioned media, 10 ng/ml VEGF (R&D systems: 293-VE-010) or 10 μg/ml SERPINE1 (R&D systems: 1786-PI-010) were subcutaneously inserted under the skin and then sutured to cover the incisions. Angioreactors were retrieved after 14 days of incubation for careful collection of the mouse endothelial cells that were attracted and invaded the cylinders. Neovascularization was quantified by staining the endothelial cells with FITC-lectin and measuring the intensity of fluorescence within a Thermo-Fisher Fluoroskan Ascent FL fluorimeter at 485nm.

### Doxycycline inducible knockdown

We employed Dharmacon SMARTvector inducible lentiviral shRNA that utilize the Tet-On 3G bipartite induction system and TurboGFP as a fluorescent reporter. This tightly regulated system consists of an inducible RNA polymerase II promoter, Phosphoglycerate kinase (PGK), which has been optimized for both minimal basal expression and potent activation upon induction by doxycycline on the TRE3G. Together, the Tet-On 3G protein and TRE3G promoter permit tight regulation of the shRNA expression, including potent induction, even at the low doxycycline doses that are required for *in vivo* xenograft studies. The selected shPAX8 (Dharmacon: V3IHSPGG-8017023), shSOX17 (Dharmacon: V3IHSPGG-6371478) and Non-Targeting Control (Dharmacon: VSC6580) high-titer lentivirus were used to transduce OVCAR3, OVCAR4 and OVTOKO cells as recommended by the manufacturer’s protocol.

### Generation of Luciferase-expressing cells

Ovarian cancer cell lines (5 × 10^4^ cells per 24 well) in complete media containing 8 μg/mL Polybrene (Millipore-Sigma: TR-1003-G) were transduced with high-titer Luc-mCherry lentivirus (64). Lentivirus was produced with 3rd Generation Packaging System Mix (Abcam: LV053) in 293T cells transfected with Lipofectamine 3000 Reagent (Thermo-Fisher: L3000008) in OPTI-MEM medium (Gibco: 31985-062). In order to produce high-titer particles, 293T transfected cells conditioned media were condensed with the Lenti-X Concentrator (Clontech: 631231) at 4°C for 24 hours.

### Mice xenograft

Following IACUC review and approval (Protocol #806825), one million OVTOKO harboring shRNA-luciferase-positive cells were inoculated intraperitoneally into five-six weeks old female immunodeficient NSG mice (JAX: 005557). All NSG mice cohorts were aseptically housed under the Stem Cell and Xenograft Core barrier. After 72 hours post-injections, feeding supplemented with 1g/kg of doxycycline (Teklad: TD.06294) was made continuously available for the mice. Tumor progression was examined by bioluminescence imaging using the Lumina IVIS system and 150 mg/kg of Luciferin (Gold Biotechnology, 115144-35-9) in PBS. Animals were sacrificed when they reached IACUC pre-determined endpoints for ascites and tumor collection.

### Statistical analysis

Representative graphics are displayed as mean and standard derivation of experiments replicates. Significant changes *P* < 0.05 between controls and knockdowns were acquired applying Student’s test (GraphPad Prims 8).

## ACKNOWLEDGMENTS

We recognize the Drapkin laboratory members for their helpful consideration and support in especially Kai Doberstein. We also acknowledge Cynthia Lopez-Haber for her technical insights, Gregory Dressler for sharing the PAX8-Luciferase reporter vector, and Andrew Kung for donating the FUW-Luc-mCherry vector. In addition, we are very grateful for the Stem Cell & Xenograft Core, the Quantitative Proteomics Resource Core, and the MD Anderson RPPA Core (NCI P30 CA16672) for technical assistance and resource access. We are also very appreciative of Tony Secreto and Lauren Schwartz support and expertise.

This work was supported by an NIH SPORE in ovarian cancer P50 CA228991 (R.D.), the Dr. Miriam and Sheldon G. Adelson Medical Research Foundation (R.D.), the Honorable Tina Brozman Foundation for Ovarian Cancer Research (R.D.), the Basser Center for BRCA (R.D.), the Claneil Foundation (R.D.), an NIH P01CA196539 (B.A.G), the Ann and Sol Schreiber Mentored Investigator Award (545754) from the Ovarian Cancer Research Alliance (D. C-M), and the Liz Tilberis Early Career Award (599175), also from the Ovarian Cancer Research Alliance (K.L.).

## SUPPLEMENTAL INFORMATION

**SI 1:**
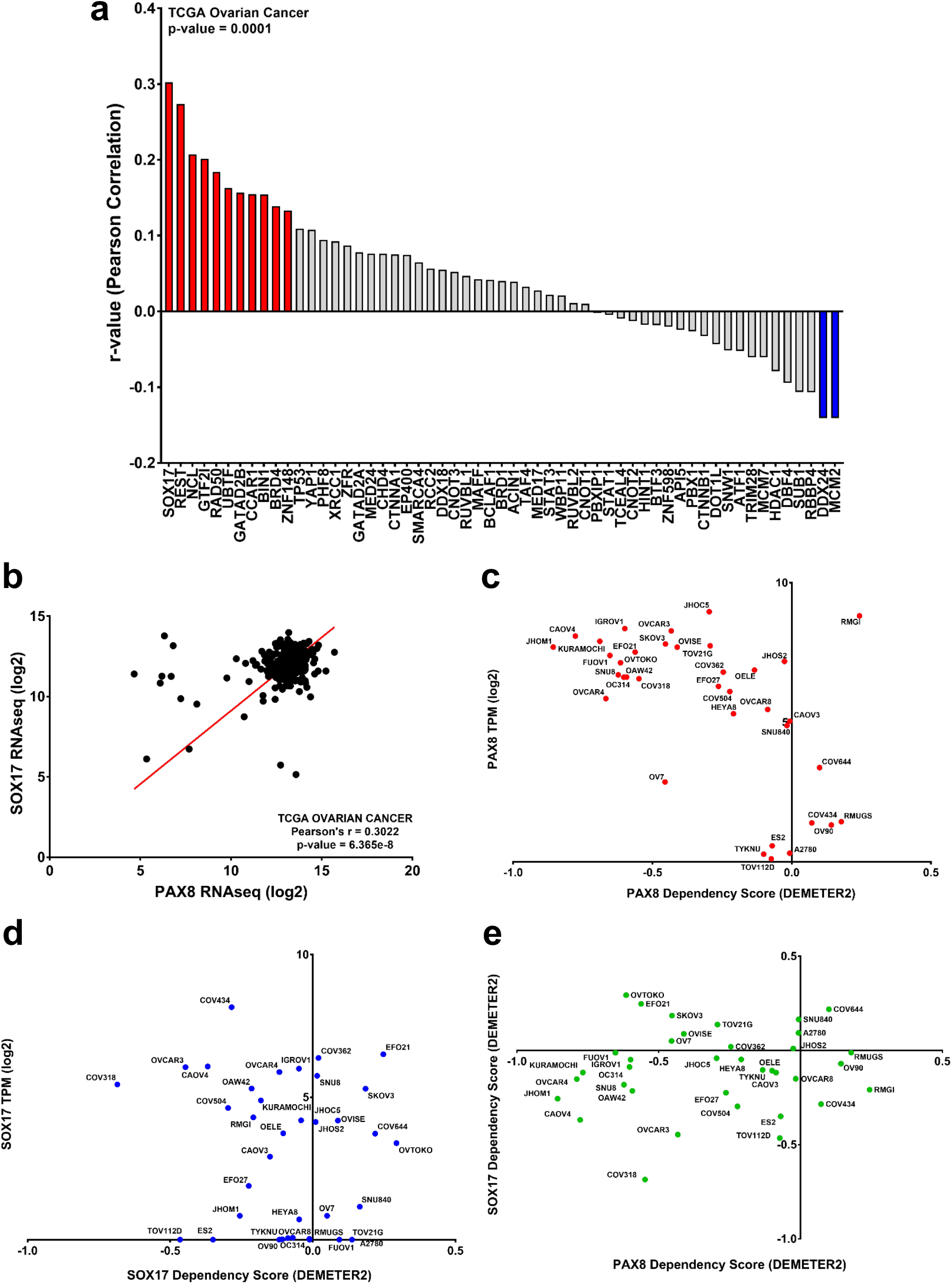
PAX8 and SOX17 have a strong correlation and dependency on ovarian cancer. (a) TCGA ovarian cancer Pearson correlation of PAX8 with the newly identified PAX8-interacting partners. (b) SOX17 has the strongest correlation with PAX8 in ovarian cancer. (c) Ovarian carcinoma cells PAX8 dependency. (d) Ovarian carcinoma cells SOX17 dependency and (e) Ovarian carcinoma cells PAX8-SOX17 co-dependency.

**SI 2:**
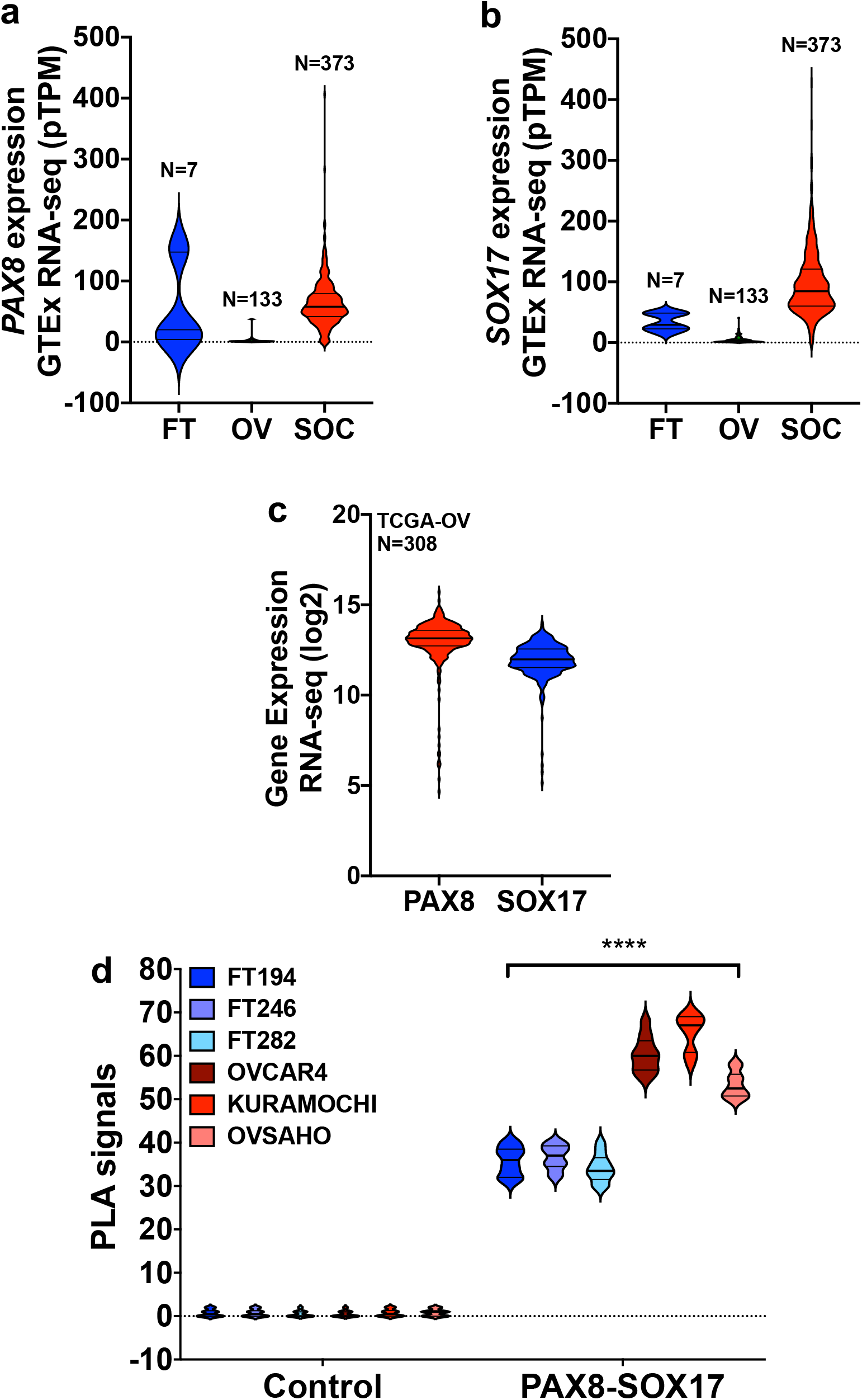
Overexpression of PAX8 and SOX17 in ovarian carcinoma cases. (a) Higher expression of PAX8 in ovarian cancer samples. (b) Differential expression of SOX17 in ovarian cancer samples. (c) Most of the TCGA ovarian cancer samples have high PAX8 and SOX17 co-expression. (d) Higher number of PAX8-SOX17 interactions on ovarian cancer cell lines.

**SI 3:**
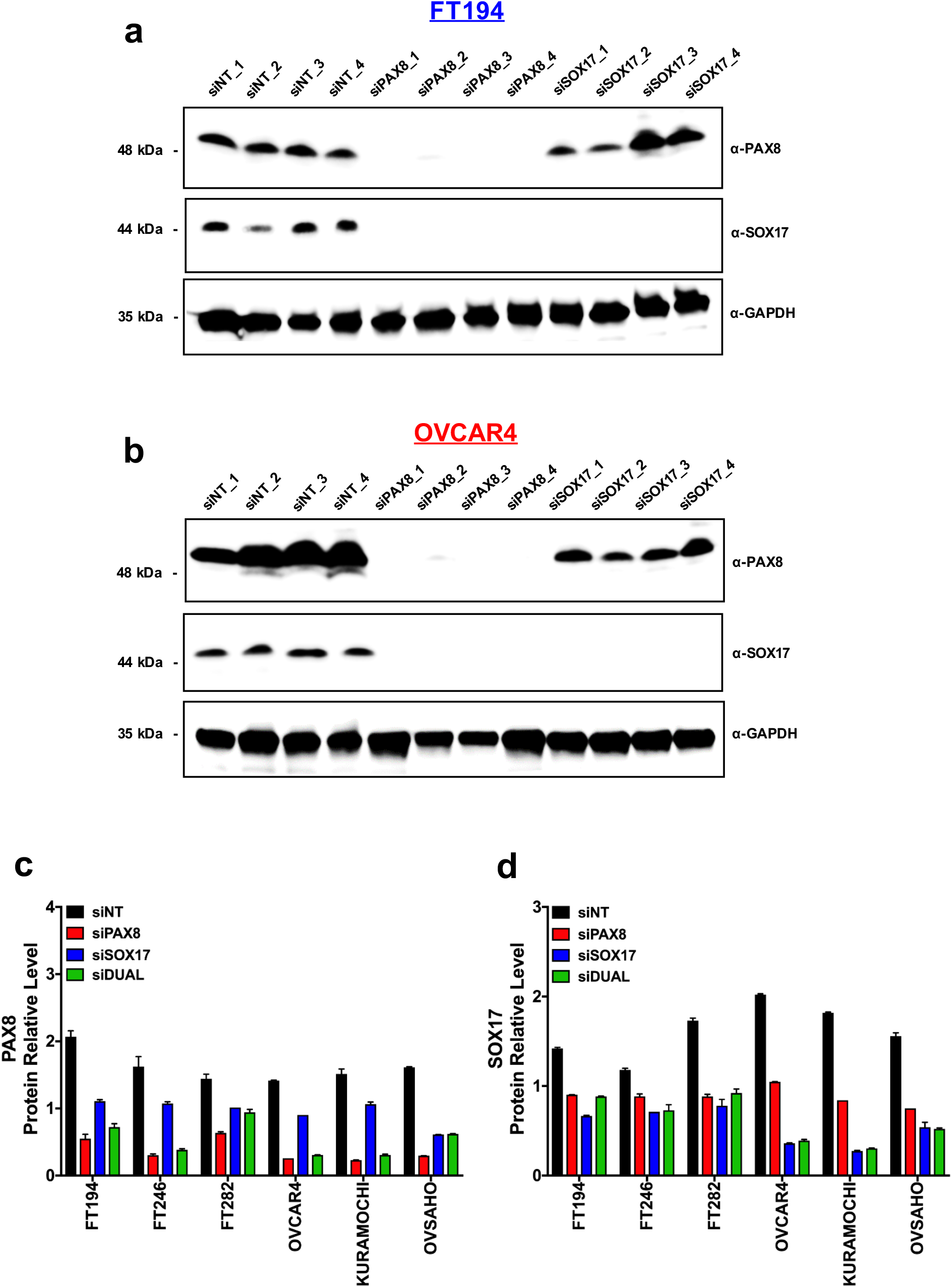
PAX8-SOX17 expression co-regulation. (a-b) Immunoblot of four deconvoluted siPAX8 and four deconvoluted siSOX17 knockdowns on FTSEC and HGSOC cell lines. (c-d) PAX8 and SOX17 protein levels after knockdowns with four pooled siPAX8, four pooled siSOX17 or then all combined by reverse-phase protein array.

**SI 4:**
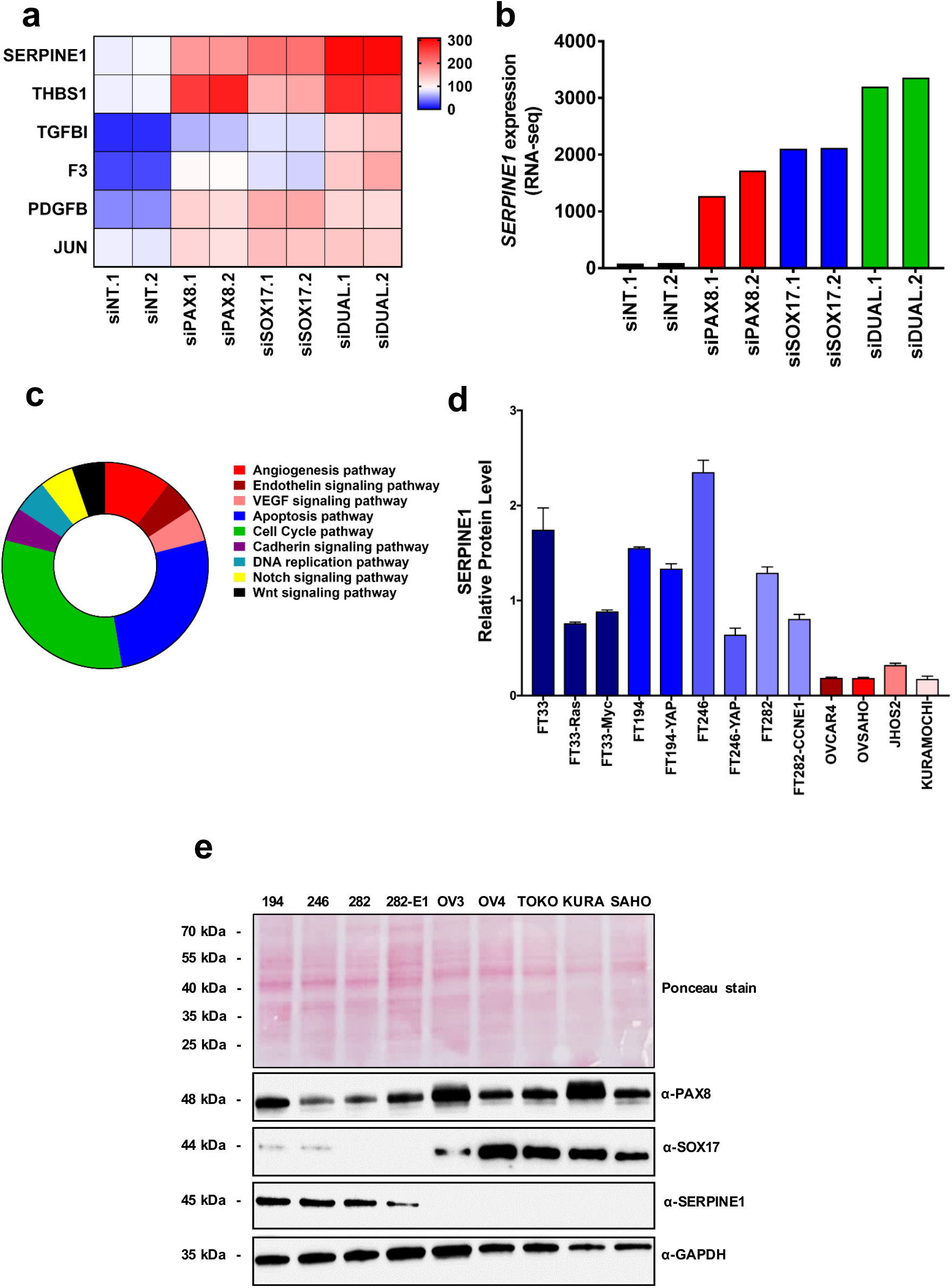
PAX8-SOX17 tightly suppresses SERPINE1 expression. (a) RNA-seq analysis depicting *SERPINE1* gene up-regulation after PAX8, SOX17 or DUAL knockdown. (b) *SERPINE1* average expression after PAX8, SOX17 or DUAL knockdown. (c) RPPA ontology analysis corroborating enrichment of angiogenesis and VEGF pathways. (d) SERPINE1 relative protein levels in different FTSEC lines and HGSOC lines, depicting suppression of SERPINE1 during malignant transformation. (e) Immunoblot of different benign and malignant cell lines showing drastically reduction of SERPINE1 levels in cell lines with increased SOX17 expression.

**SI 5:**
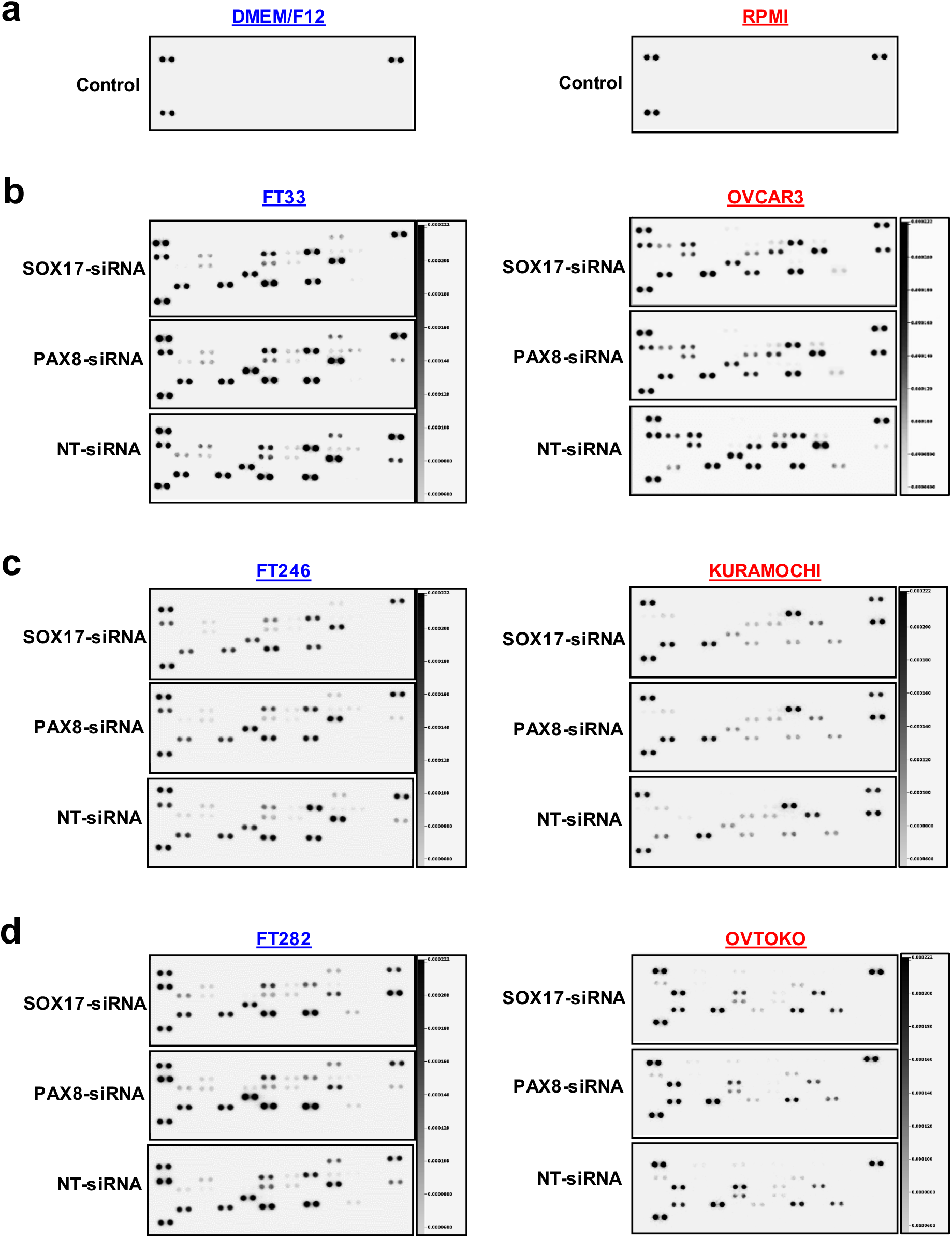
PAX8 and SOX17 regulate the secretion of angiogenesis mediators. (a) Human angiogenesis arrays of fresh DMEM/F12 and RMPI media with no detection of angiogenesis mediators, negative controls. (b-c-d) Human angiogenesis array of additional three fallopian tube secretory cells and additional three ovarian carcinoma cells conditioned media after PAX8 or SOX17 knockdown.

**SI 6:**
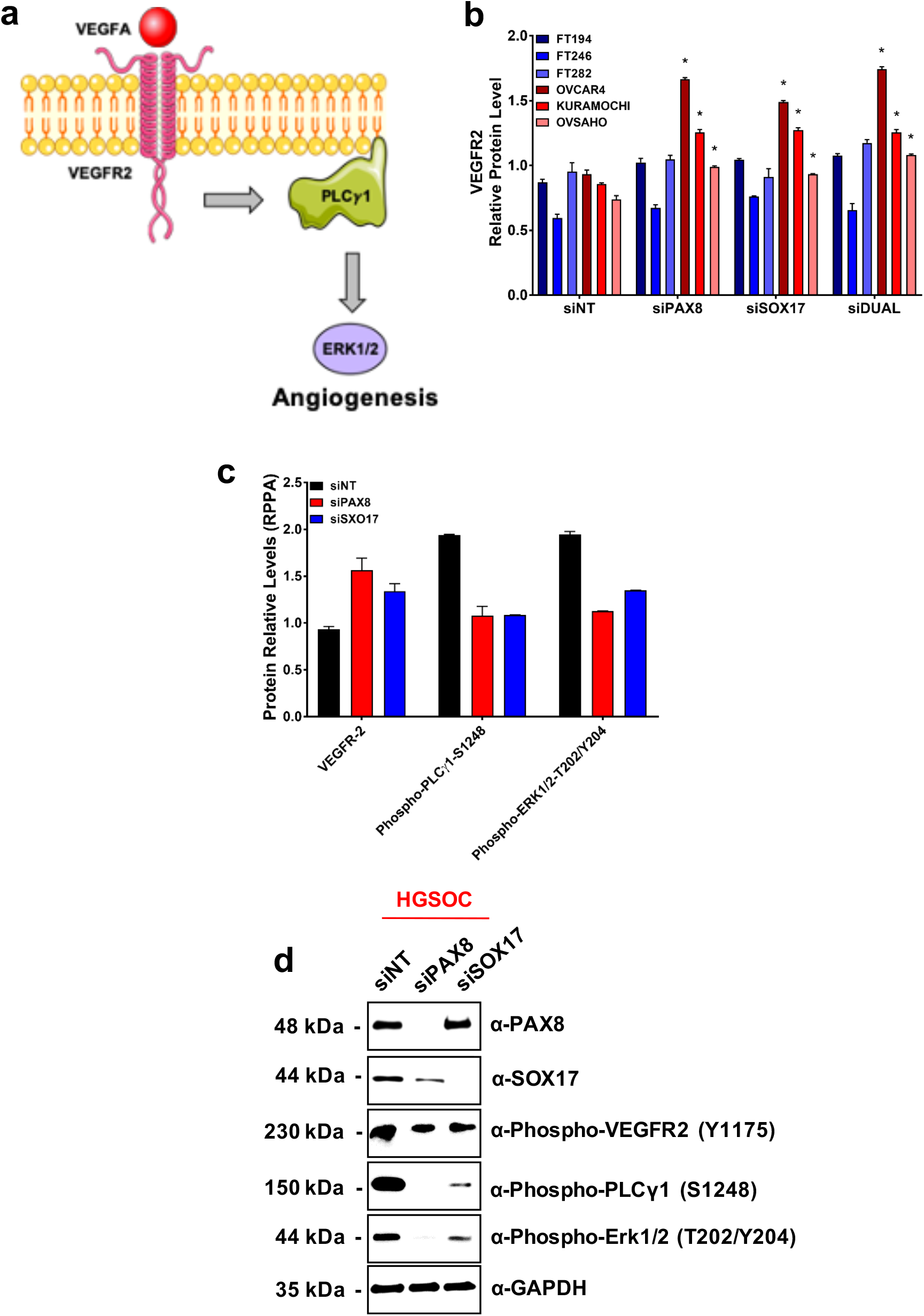
PAX8-SOX17 regulates angiogenesis through the PLCy1 pathway. (a) Schematic illustration of the PLCy1 pathway. (b) RPPA analysis showing up-regulation of VEGFR2 protein levels followed PAX8 and SOX17 knockdown. (c) RPPA analysis showing reduced phosphorylated-PLCy1 and phosphorylated-ERK1/2 (active molecules) followed PAX8 and SOX17 knockdown. (d) Confirmation by western blot of the PLCy1 pathway inactivation.

**SI 7:**
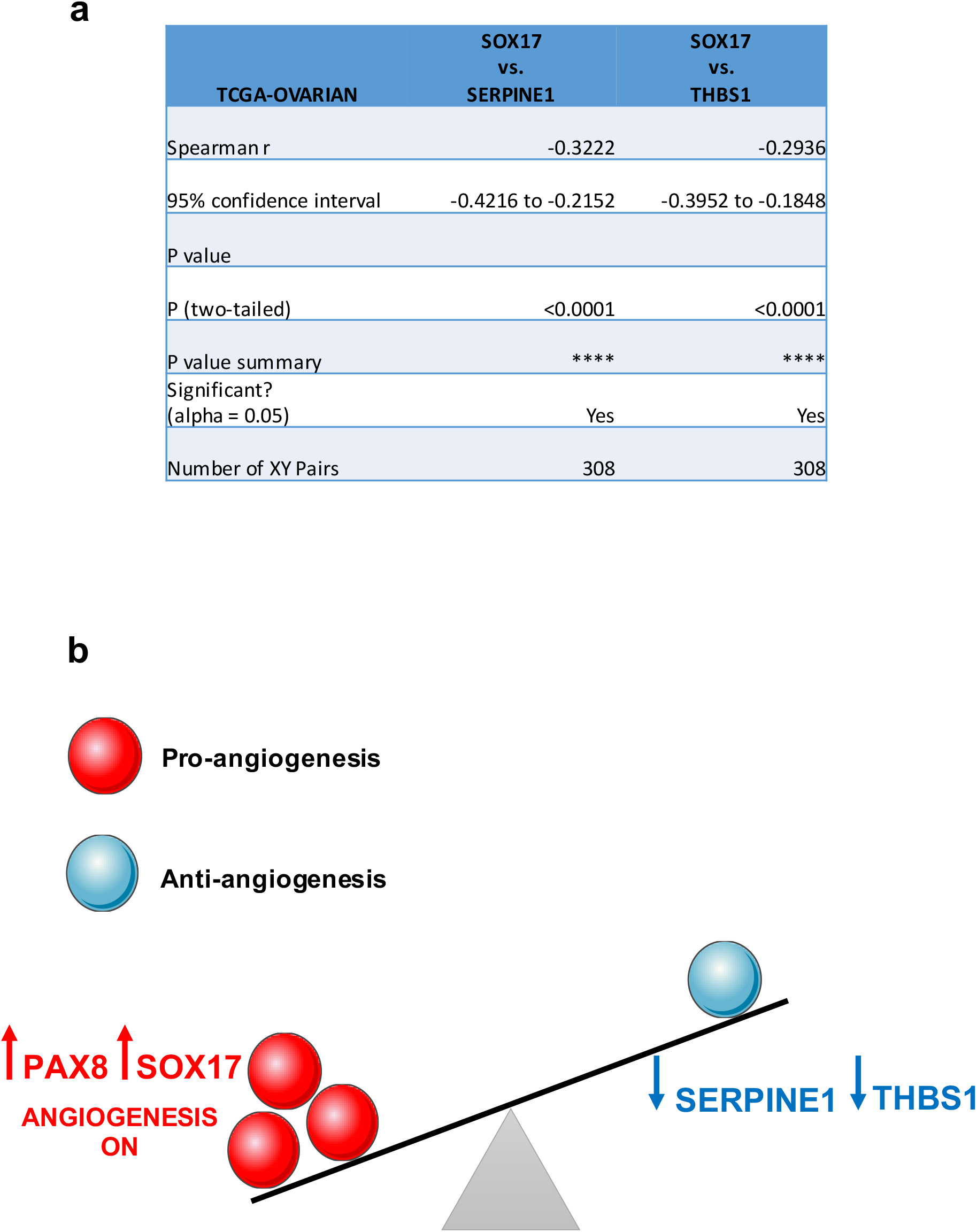
PAX8-SOX17 suppress anti-angiogenesis factors. (a) Spearman correlation analysis of TCGA ovarian cancer data set depicting negative inversely correlation between SOX17 and angiogenesis inhibitors (SERPINE1 and THBS1). (b) Schematic diagram of the activation of tumor angiogenesis by PAX8 and SOX17 expression.

## Notes

### Competing Interest Statement

R. Drapkin is a consultant/advisory board member for Repare Therapeutics and Siamab Therapeutics. G. Konecny receives honoraria from Clovis, AZ, and GSK/Tesaro and research support from Merck and Eli-Lilly. No other authors declare potential conflicts of interest.

